# Introgression between highly divergent fungal sister species

**DOI:** 10.1101/2022.08.26.505392

**Authors:** Vilde Bruhn Kinneberg, Dabao Sun Lü, David Peris, Mark Ravinet, Inger Skrede

## Abstract

To understand how species evolve and adapt to changing environments, it is important to study gene flow and introgression due to their influence on speciation and radiation events. Here, we apply a novel experimental system for investigating these mechanisms using natural populations. The system is based on two fungal sister species with morphological and ecological similarities occurring in overlapping habitats. We examined introgression between these species by conducting whole genome sequencing of individuals from populations in North America and Europe. We assessed genome wide nucleotide divergence and performed crossing experiments to study reproductive barriers. We further used ABBA-BABA statistics together with a network analysis to investigate introgression, and conducted demographic modelling to gain insight into divergence times and introgression events. The results revealed that the species are highly divergent and incompatible in vitro. Despite this, small regions of introgression were scattered throughout the genomes and one introgression event likely involves a ghost population (extant or extinct). This study demonstrates that introgression can be found among divergent species and that population histories can be studied without collections of all the populations involved. Moreover, the experimental system is shown to be a useful tool for research on reproductive isolation in natural populations.

## Introduction

Speciation can occur rapidly, changing the course of evolution in a single event, or over a long period of time with gradual shifts from semi-compatible populations to complete divergence (Nosil et al. 2017). When speciation occurs gradually, barriers to gene exchange do not arise immediately and gene flow between diverging populations can be maintained. Consequently, hybrid individuals may form (Harrison and Larson 2014; Ravinet et al. 2018). If these hybrids backcross into one of the parental species, a scenario termed introgression (Anderson and Hubricht 1938; Aguillon 2022), it can result in unique genetic combinations (Stukenbrock 2016). The amalgamation of genes across lineages can also contribute to the strengthening of barriers to gene exchange. These barriers can arise when selection increases reproductive isolation, a process known as reinforcement (Butlin 1987). Hybrids can contribute to reinforcement because they often have detrimental gene combinations, resulting in poorer fitness compared to the parental species (Abbott et al. 2013). Consequently, hybridization can both give rise to beneficial gene combinations which selection can act upon, and at the same time accelerate the divergence process by contributing to reproductive barriers (Abbott et al. 2010).

Hybridization is established as a common event in nature and can lead to the formation of hybrid species (Mallet et al. 2016; Ackermann et al. 2019; Eberlein et al. 2019; Grant and Grant 2019), as seen in several taxa including plants (e.g., *Senecio* spp.; Hegarty and Hiscock 2005; Wood et al. 2009), animals (e.g., *Heliconius* butterflies; Mavárez et al. 2006 and *Passer italiae*; Hermansen et al. 2011), and certain fungal groups (e.g., *Saccharomyces* spp.; Langdon et al. 2019 and *Zymoseptoria pseudotritici*; Stukenbrock et al. 2012). Even though hybridization seems to be common in contemporary populations, we do not know the impact it will have on future populations. By studying the genomes of contemporary species with a history of hybridization and introgression, it may be possible to understand how previous interspecific gene exchange have influenced the populations we observe today and infer the future effect of current events.

When taxa have diverged over a long period of time, it can be difficult to discover ancient admixture as genomic signals of introgression can be blurred over macroevolutionary time. Moreover, detection might be difficult due to the deleterious nature of most introgressed genes between divergent species, or lack of time for introgressed regions to spread in the population if the gene flow is recent (Maxwell et al. 2018). Hence, the evolutionary history of a genus can be complex even though current investigations recover clear and resolved phylogenies (Keuler et al. 2020).

Signs of ancient or low frequency introgression have been possible to detect using high-throughput sequencing and statistical models (e.g., Crowl et al. 2019; Ravinet et al. 2018). Regions of introgression might constitute small parts of otherwise divergent genomes due to erosion of linkage by recombination coupled with a long period of mostly independent evolution or purifying selection (Maxwell et al. 2018; Ravinet et al. 2018; Schumer et al. 2018; Martin et al. 2019; Cuevas et al. 2022). The retention of specific introgressed regions can for example represent adaptational benefits (Racimo et al. 2015), regions of high recombination rate (Nachman and Payseur 2012; Ravinet et al. 2018; Schumer et al. 2018), and regions under lower constraint or less purifying selection (Schumer et al. 2016). However, the patterns of introgression can also be difficult to distinguish from mechanisms such as incomplete lineage sorting (i.e., preservation of ancestral polymorphisms; Platt et al. 2019). Methods have been developed to circumvent confounding signals (e.g., ABBA-BABA statistics; Green et al. 2010; Durand et al. 2011) and in principle it is possible to separate introgression from other evolutionary processes (Martin et al. 2014). Research on introgression between divergent species can reveal important contributions to the evolutionary history of the taxa involved (e.g., ecological adaptations; Nelson et al. 2021) and increase our understanding of how such mechanisms can affect contemporary populations in the future and how robust reproductive barriers are against gene flow between divergent species.

There is currently a need for tractable experimental systems to study reproductive isolation in natural populations (White et al. 2019; Stankowski and Ravinet 2021). An interesting experimental system for investigating such processes appear in fungal species complexes in the Agaricomycotina. The subphylum Agaricomycotina is a diverse taxon, with about 20,000 species described worldwide and a crown age estimate of around 429 million years (Floudas et al. 2012). Research on hybridization and introgression among species of the Agaricomycotina (mushroom-forming fungi) is limited, but there are some examples from genera including *Pleurotus* (Bresinsky et al. 1987), *Heterobasidion* (Garbelotto et al. 1996; Stenlid and Karlsson 1991; Giordano et al. 2018), and *Armillaria* (Baumgartner et al. 2012), indicating that hybridization may be a common but understudied mechanism of speciation and gene exchange among taxa in this branch of the tree of life. Moreover, recent research shows that the reproductive barrier in fungi can be permeable despite high divergence between species (Maxwell et al. 2018), making the fungal reproductive system an interesting case study for expanding our knowledge on reproductive isolation and the speciation continuum (Maxwell et al. 2018).

In this study we use natural populations of the sister species *Trichaptum fuscoviolaceum* (Ehrenb.) Ryvarden and *Trichaptum abietinum* (Dicks.) Ryvarden (pictured in Figure 2) as our model organisms. These fungi are saprotrophic white rot fungi growing on conifers across the northern hemisphere. The two species are broadly sympatric and can grow on the same host, and sometimes they are found together on the same substrate. They are in general phylogenetically well separated species (Kauserud and Schumacher 2003; Seierstad et al. 2020; Peris et al. 2022), but some individuals clustered incongruently for different loci in a previous study including only a few molecular markers (Seierstad et al. 2020). Here, the authors suggested introgression or incomplete lineage sorting as possible explanations for the conflicting phylogenetic signals (Seierstad et al. 2020). Reproductive barriers between the two species have been documented by in vitro crossing experiments (Macrae 1967).

Population structure has been found within *T. abietinum* (Seierstad et al. 2020; Peris et al. 2022). In North America, two populations referred to as the North American A and the North American B population occur in sympatry and are reproductively isolated (i.e., form intersterile groups; Macrae 1967; Magasi 1976; Peris et al. 2022). Such intersterile groups have not been detected among populations of *T. fuscoviolaceum* (Macrae 1967; Peris et al. 2022), but some genetic structuring of populations has been observed (Seierstad et al. 2020; Peris et al. 2022).

Through this study, we aimed at exploring how signs of introgression can be discovered in genomes of extant species and how regions retained from past or current introgression might influence the evolution of contemporary populations. We used the experimental system to investigate potential introgression by conducting whole genome sequencing of individuals belonging to different populations of *T. abietinum* and *T. fuscoviolaceum*. Further, we assessed the possibility of current gene flow by testing compatibility across species through in vitro crossing experiments. We hypothesised that *T. fuscoviolaceum* and *T. abietinum* populations do not hybridize frequently due to a well resolved phylogeny and earlier crossing experiments revealing the species to be intersterile (Macrae 1967; Seierstad et al. 2020; Peris et al 2022). However, due to their overlapping habitat, ecology, and morphology, we hypothesised that the sister species might have a shared history that involves introgression, possibly having occurred among ancient populations.

## Materials and Methods

### Sampling

Individuals of *T. abietinum* and *T. fuscoviolaceum* were collected in New Brunswick, Canada and Pavia, Italy during the autumn of 2018. One individual of *T. biforme* was collected in New Brunswick, Canada, and included as an outgroup. For all collection sites, ten individuals were sampled from separate logs, or two meters apart on the same log, within one square kilometre. For every individual, a cluster of sporocarps (covering no more than 3 × 3 cm) was collected and placed in separate paper bags. Notes on host substrate, GPS coordinates and locality were made for all individuals, and they were given a collection ID according to collection site and species identification based on morphology. The sporocarps were dried at room temperature for 2 – 3 days, or in a dehydrator at 30 °C, and later stored at room temperature in paper bags. Individuals included in this study are presented in Table S1.

### Culturing

Since haploid sequences are bioinformatically convenient to work with, we isolated monokaryotic mycelia for sequencing by culturing collected *Trichaptum* individuals as follows (illustrated in Figure 1): (A) To revive dried individuals for spore shooting, sporocarps were placed in a moist paper towel and left in the fridge until soaked through (about 3 hours). (B) While working in a safety cabinet (Labculture^®^ ESCO Class II Type A2 BSC, Esco Micro Pte. Ltd., Singapore), sporocarps were attached, hymenium facing media, with silicon grease from Merck Millipore (Darmstadt, Germany) to the lid of a petri dish containing 3% malt extract agar (MEA), with antibiotics and fungicides (10 mg/l Tetracycline, 100 mg/l Ampicillin, 25 mg/l Streptomycin and 1 mg/l Benomyl) to avoid contamination. The sporocarps were left for a minimum of one hour for spores to shoot onto the MEA plates. Subsequently, the sporocarps were removed to minimize spore shooting and the petri dish was sealed off with Parafilm M^®^ (Neenah, WI, USA). The cultures were left for approximately one week, or until hyphal patches were visible, at 20 °C in a dark incubator (Termaks AS KB8400/KB8400L, Bergen, Norway). (C) Working in a safety cabinet, single, germinated spores were picked with a sterile scalpel and placed onto new MEA dishes with antibiotics and fungicides. The new cultures were left in a dark incubator at 20 °C for a few days until a mycelial patch could be observed. (D) The hyphae were checked for clamp connections in a Nikon Eclipse 50i light microscope (Tokyo, Japan) using 0.1% Cotton Blue to accentuate cells (examples in Figure S5). Clamps indicate a dikaryotic hyphae and we proceeded with the cultures lacking clamps (i.e., monokaryotic hyphae). (E) Monokaryotic cultures were replated onto new MEA dishes without antibiotics and fungicides (the mycelia grow better without these substances and the cultures were now free from contaminants) and placed in an incubator at 20 °C prior to sequencing and experiments.

**Figure 1.**
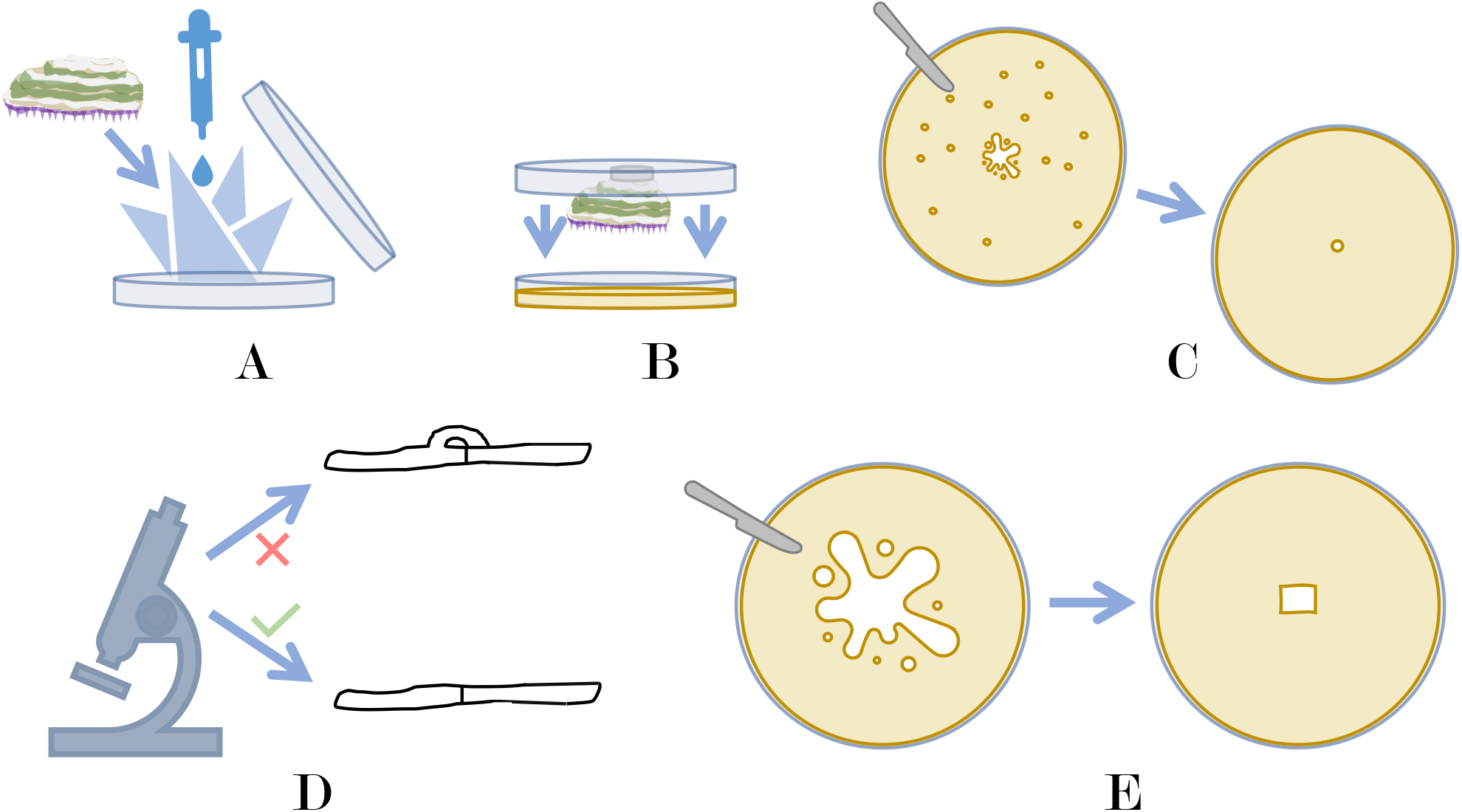
Procedure for culturing monokaryotic fungal individuals. (A) A sporocarp is placed in a wet paper towel onto a petri dish. (B) The sporocarp is glued to the lid of the petri dish to allow for spore shooting onto agar. (C) Hyphae from single spores are picked with a scalpel and placed onto new agar. (D) Microscopy of hyphae to confirm monokaryotic cultures. Hypha with clamp connection is indicated with a red cross and hypha without clamp connection is indicated with a green check symbol. (E) Mycelium from the new culture made in (C) is cut out with a scalpel and placed onto new agar.

### PCR and Sanger sequencing

We Sanger sequenced the internal transcribed spacer (ITS; the fungal barcode), to confirm correct species designation of the cultures. The ITS region was amplified using the ITS1 (5’ – TCCGTAGGTGAACCTGCGG – 3’; White et al. 1990) and ITS4 (5’ – TCCTCCGCTTATTGATATGC – 3’; White et al. 1990) primers and the Thermo ScientificTM Phire Plant Direct PCR Kit (Waltham, USA) according to the manufacturer’s protocol (using a small piece of mycelia instead of plant tissue). The following PCR program was used: 4 min at 95 °C, followed by 40 cycles of 25 sec at 95 °C, 30 sec at 53 °C and 60 sec at 72 °C, followed by a 10 min extension at 72 °C and an indefinite hold at 10 °C. PCR products were purified using 0.2 μl ExoProStar 1-Step (GE Healthcare, Chicago, USA), 1.8 μl H_2_O and 8 μl PCR product. The samples were Sanger sequenced by Eurofins Scientific (Hamburg, Germany).

We assessed, trimmed, and aligned the resulting forward and reverse sequences to consensus sequences using Geneious Prime v2020.1.2 (https://www.geneious.com). To verify species designation of cultures, the consensus sequences were checked with the Basic local alignment search tool (BLAST; Altschul et al. 1990) against the National Centre of Biotechnology Information (NCBI) database (U.S. National Library of Medicine, Bethesda, MD, USA). We kept cultures identified as *T. fuscoviolaceum* or *T. abietinum* and updated the *Trichaptum* cultures with incorrect initial species designation (based on sporocarp morphology).

### DNA extraction and Illumina sequencing

Where possible, we chose approximately five individuals of *T. fuscoviolaceum* from each collection site and five individuals of *T. abietinum* corresponding to the same sites for DNA extraction and Illumina sequencing, including one individual of *T. biforme* as an outgroup.

Mycelia from fresh cultures, grown on 3% MEA with a nylon sheet between the MEA and the mycelia, were scraped off the plate and DNA was extracted using the E.Z.N.A.^®^ Fungal HP DNA Kit (Omega Bio-Tek, Norcross, GA, USA) according to the DNA extraction procedure explained in Peris et al. (2022). Illumina libraries were prepared by the Norwegian Sequencing Center (NSC) as explained in Peris et al. (2022). Samples were sequenced at NSC on either the Illumina Hiseq 4000 or the Illumina Novaseq I. The samples where distributed across sequencing runs (3 runs).

### Crossing experiments

To assess mating compatibility between species (i.e., between individuals of *T. abietinum* and *T. fuscoviolaceum*), we performed crossing experiments. As a positive control, we also crossed individuals of *T. fuscoviolaceum*. Some of the crosses include European *T. abietinum* that are not sequenced in the present study, but included in Peris et al. (2022). The crossing set-ups were planned according to a mating compatibility scheme based on mating loci (*MAT*) predicted in Peris et al. (2022). We crossed individuals that were both expected and not expected to mate based on their predicted mating type (i.e., dissimilar or similar allelic classes on both *MAT* loci). Individuals used for the experiment are presented in Table S2 (see Peris et al. (2022) for further details on the European *T. abietinum* individuals). Three replicates were made for all crosses to strengthen the confidence in the observations.

Pairs of monokaryotic individuals (circular 0.8 cm in diameter plugs) were plated 4 cm apart on petri dishes containing 3% MEA. The petri dishes were placed in a dark incubator at 19 °C until the two mycelia had grown together (about 2 weeks). The cultures were photographed using a Nikon D600 Digital Camera (Tokyo, Japan). To investigate if the crossing experiments were successful, we assessed the presence or absence of clamp connections using a Zeiss Axioplan 2 imaging light microscope (Güttingen, Germany) with Zeiss AxioCam HRc (Güttingen, Germany). The process is similar to the description in Figure 1D. Microscopic photographs of hyphae were taken at 400 and 630 × magnification.

### Reference genomes

We used the two genomes of *T. abietinum* (strain TA10106M1) and *T. fuscoviolaceum* (strain TF100210M3) as reference genomes (Bioproject PRJNA679164; https://doi.org/10.5061/dryad.fxpnvx0t4; Peris et al. 2022). In addition, we made a combined reference genome by merging the *T. abietinum* (acc. no. GCA 910574555) and *T. fuscoviolaceum* (acc. no. GCA 910574455) reference genomes with *sppIDer* (Langdon et al. 2018).

### Preparation and initial mapping of whole genome data

Illumina raw sequences were quality filtered, removing sequences with a Phred quality score less than 30, using *Trim Galore! v0.6.2* (https://www.bioinformatics.babraham.ac.uk/projects/trim_galore/; Krueger 2015), and assessed using *FastQC* (Andrews 2010) and *MultiQC* (Ewels et al. 2016). After pre-processing, we used *BWA v0.7.17* (Li and Durbin 2009) to search for recent hybrids by mapping Illumina reads from *T. fuscoviolaceum* to a combined reference genome of *T. fuscoviolaceum* and *T. abietinum* using the wrapper *sppIDer* (Langdon et al. 2018). The wrapper generates a reference genome with chromosomes from both *T. fuscoviolaceum* and *T. abietinum*. The reads from a strain from one species can then be mapped to the combined genome. If the reads of an individual map equally well to chromosomes of both species, it indicates that the strain is a hybrid. No hybrids were revealed among the *T. fuscoviolaceum* individuals in the *sppIDer* analysis (Figure S1). All *T. fuscoviolaceum* individuals mapped with greater depth to the *T. fuscoviolaceum* part of the combined reference genome than the *T. abietinum* part. Since there were no recent hybrid individuals and we could only use one reference for further analyses, we chose to continue with the *T. fuscoviolaceum* reference genome.

### Re-mapping with Stampy

To improve mapping of *T. abietinum, T. biforme* and the Italian *T. fuscoviolaceum* to the reference genome (based on a Canadian *T. fuscoviolaceum* individual), the raw sequences were mapped with *Stampy v1.0.32* (Lunter and Goodson 2011), which is designed to be more sensitive to divergent sequences (Lunter and Goodson 2011), before continuing with further analyses. Based on nucleotide divergence estimates found in Peris et al. (2022) by conversion of average nucleotide identity using *FastANI* (Jain et al. 2018), the substitution rate flag was set to 0.23 for *T. biforme*, 0.067 for the Italian *T. fuscoviolaceum*, and 0.157 for *T. abietinum* when mapping each to the reference. The raw sequences were not trimmed before mapping due to limitations on hard clipping in *Stampy* (i.e., sequences are sometimes too short for *Stampy*), but poor sequences were filtered away at a later stage (see below).

### SNP calling and filtering

To obtain a dataset with single nucleotide polymorphisms (SNPs), we first used *GATK HaplotypeCaller v4.1.4*. (McKenna et al. 2010). To create the dictionary files and regroup the mapped files before SNP calling, we used *Picard v2.21.1* (https://broadinstitute.github.io/picard/) and reference index files were made using *SAMtools* faidx (Li et al. 2009). We ran *HaplotypeCaller* in haploid mode with otherwise default settings. Subsequently, we used the resulting Variant Call Format (VCF) files in *GATK GenomicsDBImport* (McKenna et al. 2010) to create a database used as input for *GATK GenotypeGVCF* (McKenna et al. 2010), which creates a VCF file containing SNPs for all individuals. *GenomicsDBImport* was used with default settings together with the java options (‘--java-options’) ‘-Xmx4g’ and ‘-Xms4g’ and an interval text file (‘--intervals’) containing names of the different scaffolds. *GenotypeGVCF* was used with default settings. To remove indels, bad SNPs, and individuals with high missingness, we filtered the resulting VCF file with *GATK VariantFiltration* (McKenna et al. 2010) and *BCFtools v1.9* filter (Danecek et al. 2021). We used GATK’s hard filtering recommendations (https://gatk.broadinstitute.org/hc/enus/articles/360035890471-Hard-filtering-germline-short-variants) together with the Phred quality score option of removing SNPs with a score less than 30.0 (‘QUAL < 30.0’). With *BCFtools* filter, we removed indels and poor SNPs using these options: minimum read depth (DP) < 3, genotype quality (GP) < 3 and ‘-v snps’. We also used *BCFtools* filter to remove multiallelic SNPs (‘view -M2’), SNPs close to indels (‘--SnpGap 10’), variants with a high number of missing genotypes (‘-e ‘F_MISSING > 0.2’’), minimum allele frequency (‘MAF <= 0.05’), and invariant sites and monomorphic SNPs (‘-e ‘AC==0 || AC==AN’’). We made one dataset where monomorphic SNPs were removed and the MAF filter was applied (Dataset 1 with 2 040 885 SNPs) and two datasets, one with the outgroup and one without, not applying these filters (Dataset 2 with 3 065 109 SNPs; Dataset-O 2 with 3 118 957 SNPs, where O = outgroup), because monomorphic sites were required to calculate some divergence statistics. After filtering, individuals with high missingness or high heterozygosity (i.e., dikaryons) were removed. The final datasets consisted of 32 individuals from the Canadian *T. fuscoviolaceum* population, 9 individuals from the Italian *T. fuscoviolaceum* population, 30 individuals from the North American B *T. abietinum* population, and 6 individuals from the North American A *T. abietinum* population.

### Phylogenetic tree analysis

To confirm the phylogenetic relationship between the different populations of *T. abietinum* and *T. fuscoviolaceum*, we performed a maximum likelihood phylogenetic tree analysis using *IQ-TREE 2* (Minh et al. 2020). The VCF-file from Dataset-O 2 was converted into a PHYLIP file by using Edgardo M. Ortiz’s script *vcf2phylip.py* (https://raw.githubusercontent.com/edgardomortiz/vcf2phylip/master/vcf2phylip.py). The IQ-TREE analysis was run on the PHYLIP file using the flags ‘-T auto’, ‘-m GTR+ASC’, ‘-alrt 1000’ and ‘-B 1000’. GTR+ASC is a standard model.

### Principal component and divergence analyses

To explore the data and investigate population groupings, we performed a principal component analysis (PCA) with *PLINK v2.00-alpha* (www.cog-genomics.org/plink/2.0/; Chang et al. 2015). To prepare the input file, we linkage pruned Dataset 1 in *PLINK*, using the flags ‘—vcf $vcf_file’, ‘--double-id’, ‘--allow-extra-chr’, ‘--set-missing-var-ids @:#’, --out $out_file’ and ‘--indep-pairwise 50 10 0.1’, retaining 56 046 SNPs. The ‘--indep-pairwise’ flag performs the linkage pruning, where ‘50’ denotes a 50 Kb window, ‘10’ sets the window step size to 10 bp, and ‘0.1’ denotes the r^2^ (or linkage) threshold. A PCA was subsequently performed on the pruned VCF file, using the flags, ‘--vcf $vcf_file’, ‘--double-id’, ‘--allow-extra-chr’, ‘--set-missing-var-ids @:#’, ‘--extract $prune.in_file’, ‘--make-bed’, ‘--pca’, and ‘--out $out_file’ (both linkage pruning and PCA flags were based on the Physalia tutorial https://speciationgenomics.github.io/pca/).

To investigate the divergence between populations, we applied a sliding window approach on Dataset 2 to calculate the fixation index (*F*_ST_) and absolute divergence (*d*_XY_) along the genome. We also performed a sliding window analysis to calculate within population divergence (π). The analyses were performed using Simon Martin’s script *popgenWindows.py* (https://github.com/simonhmartin/genomics_general/blob/master/popgenWindows.py) with *Python v3.8* (Van Rossum and Drake 2009). We set the window size to 20 000 bp (‘-w 20000’), step to 10 000 bp (‘-s 10000’) and the minimum number of SNPs in each window to 10 (‘-m 10’).

### Introgression analyses with D-statistics

To investigate introgression between populations, we used the *R* (R Core Team 2020) package *admixr* (Petr et al. 2019) and Dataset-O 2 to calculate the *D* (Green et al. 2010; Durand et al. 2011), outgroup *f*_3_ (Raghavan et al. 2014) and *f*_4_-ratio (Reich et al. 2009; 2011; Patterson et al. 2012) statistics between different populations (based on recommendations from the Physalia tutorial https://speciationgenomics.github.io/ADMIXTOOLS_admixr/, and the *admixr* tutorial https://bodkan.net/admixr/articles/tutorial.html#f4-ratio-statistic-1). To prepare the input file from VCF to Eigenstrat format, we used the conversion script *convertVCFtoEigenstrat.sh* by Joana Meier (https://github.com/speciationgenomics/scripts), which utilizes *VCFtools v0.1.16* (Danecek et al. 2011) and *EIGENSOFT v7.2.1* (Patterson 2006; Price 2006). The script has a default recombination rate of 2.0 cM/Mb, which we changed to 2.5 cM/Mb based on earlier findings in the class Agaricomycetes, where *Trichaptum* belongs (Heinzelmann et al. 2020). We further used another *Python* script developed by Simon Martin, *ABBABABAwindows.py* (https://github.com/simonhmartin/genomics_general/blob/master/ABBABABAwindows.py), for a sliding window ABBA-BABA analysis on Dataset-O 2 to calculate the proportion of introgression (*f*_dM_; Malinsky et al. 2015). The window size was set to 20 000 (‘-w 20000’), step size to zero, and minimum number of SNPs per window to 100 (‘-m 100’), together with ‘--minData 0.5’ to specify that at least 50% of the individuals in each population must have data for a site to be included (based on recommendations from the Physalia tutorial https://speciationgenomics.github.io/sliding_windows/). We used *T. biforme* as outgroup and tested introgression between the Canadian *T. fuscoviolaceum* and the *T. abietinum* populations in addition to the Italian *T. fuscoviolaceum* and the *T. abietinum* populations (the phylogenetic topology was based on results from the *f*_3_ analysis). Outlier windows were extracted from the results with a Hidden Markov-model approach using the *R* package *HiddenMarkov* (Harte 2021) following Ravinet et al. (2018). Since the HMM approach cannot analyse negative values, the *f*_dM_ distribution was rescaled by adding 2 to all values. Annotated genes in these outlier windows were retrieved from the annotated *T. fuscoviolaceum* reference genome. The reference genome was annotated using *RepeatModeler* (Flynn et al. 2020), *RepeatMasker* (Smit et al. 2013-2015) and *MAKER2* (Holt and Yandell 2011). Functional annotation and protein domain annotations of detected coding sequences and the encoded proteins were performed using *blastp* (Altschul et al. 1990) against a local UniProt database and InterProScan (Jones et al. 2014), respectively. All annotations were encoded in a General Feature Format (GFF) file, which was used to match the significant windows and extract the genes. Gene ontology terms annotated in the GFF file were extracted using the package *rtracklayer v1.48* (Lawrence et al. 2009) in *R*. Gene ontology (GO) enrichment analysis was performed in *R* using the *TopGO v2.40* package (Alexa and Rahnenfuhrer 2021). Lastly, a false discovery rate (FDR) analysis was performed on the resulting raw p-values.

### Network analysis with TreeMix

To further explore possible introgression events and direction of introgression, we applied a network analysis with *TreeMix* (Pickrell and Pritchard 2012). To prepare the VCF-file (Dataset-O 2) for analysis, we removed sites with missing data using *VCFtools* (with ‘--max-missing 1’) and linkage pruned the data using *PLINK*. Linkage pruning was performed in the same way as with the PCA, except the file was recoded into a new VCF file using the flags ‘--bfile’ and ‘--recode vcf’ after pruning. To convert the data to *TreeMix* format, we ran Joana Meier’s script *vcf2treemix.sh* (https://github.com/speciationgenomics/scripts/blob/master/vcf2treemix.sh). The script *vcf2treemix.sh* also requires the script *plink2treemix.py* (https://bitbucket.org/nygcresearch/treemix/downloads/plink2treemix.py).

*TreeMix* was run using the options ‘-global’ and ‘-root *T_biforme*’. The option for number of edges (‘-m’) was analysed from 1-10 and the option for block size (‘-k’) was varied between 300 and 800 to avoid identical likelihoods. For each block size, the analysis was repeated three times for ‘m’ 1-10 edges. To find the optimal number of edges, we ran *OptM* (Fitak 2021) in *R*. The residuals and network with different edges were plotted in *R* using the functions provided by *TreeMix* (*plotting_funcs.R*) together with the packages *RColorBrewer v1.1-2* (Neuwirth 2014) and *R.utils* (Bengtsson 2021).

### Demographic modelling

To explore divergence and introgression, we applied demographic modelling with *fastsimcoal2* (Excoffier et al. 2021). To prepare the data (Dataset 2) for analysis, frequency spectrum files were created using *easySFS* (https://github.com/isaacovercast/easySFS). Subsequently, *fastsimcoal2* was run using the output files from *easySFS* together with a template file defining the demographic model and a parameter estimation file. The analyses were run with the flags ‘-m’, ‘-0’, ‘-n 200000’, ‘-L 50’, ‘-s 0’ and ‘-M’. Each model was run 100 times and the run with the best likelihood was extracted using Joana Meier’s script *fsc-selectbestrun.sh* (https://raw.githubusercontent.com/speciationgenomics/scripts/master/fsc-selectbestrun.sh).

We selected several different models based on likely events inferred from the results of the ML tree, *D* and *f* statistics, and *TreeMix* analyses to test for different scenarios of introgression, both with and without one or two ghost populations (Figure S7). To compare different models, an AIC value was calculated from the run with the best likelihood for each model using the *R* script based on code by Vitor Sousa, *calculateAIC.sh* (https://github.com/speciationgenomics/scripts/blob/master/calculateAIC.sh). Since AIC can overestimate support for the best model when SNPs are in linkage, we also calculated the likelihood distributions using the best run from all the models. The models were run with the best parameter values ({PREFIX}_maxL.par output from the first run) 100 times in *fastimcoal2* with the options ‘-n 1000000’, ‘-m’, ‘-q’, and ‘-0’. The likelihood values were collected and plotted in *R* for comparison between models (i.e., look for overlapping distributions).

## Results

### Crossing experiments confirm incompatibility between species

To ensure that the previous results of intersterility between *T. abietinum* and *T. fuscoviolaceum* were also the case for our collections, we performed new crossing experiments with our individuals. We did not observe clamp connections between crosses of *T. fuscoviolaceum* and *T. abietinum* individuals. This was the case for mate pairs that were predicted to mate based on mating type alleles and for those predicted not to (Table S2; Figure S2 and S4; mating type alleles were annotated in Peris et al. 2022). It was difficult to observe compatible crosses by investigating the cultures macroscopically, but there was often a sharper line between individuals on the petri dish when the crosses were incompatible (Figure S2). The *T. fuscoviolaceum* individuals mated as expected (i.e., those that were predicted to be incompatible due to identical mating types showed no clamp connections and those that were predicted to be compatible had clamp connections; Table S2; Figure S3 and S5).

### Phylogeny, principal component and divergence analyses reveal high divergence between species

After confirming mating incompatibility, we continued with assessing the nucleotide divergence between the species. The maximum likelihood (ML) phylogeny clustered the species and populations into well-defined clades with high support (Figure 2).

**Figure 2.**
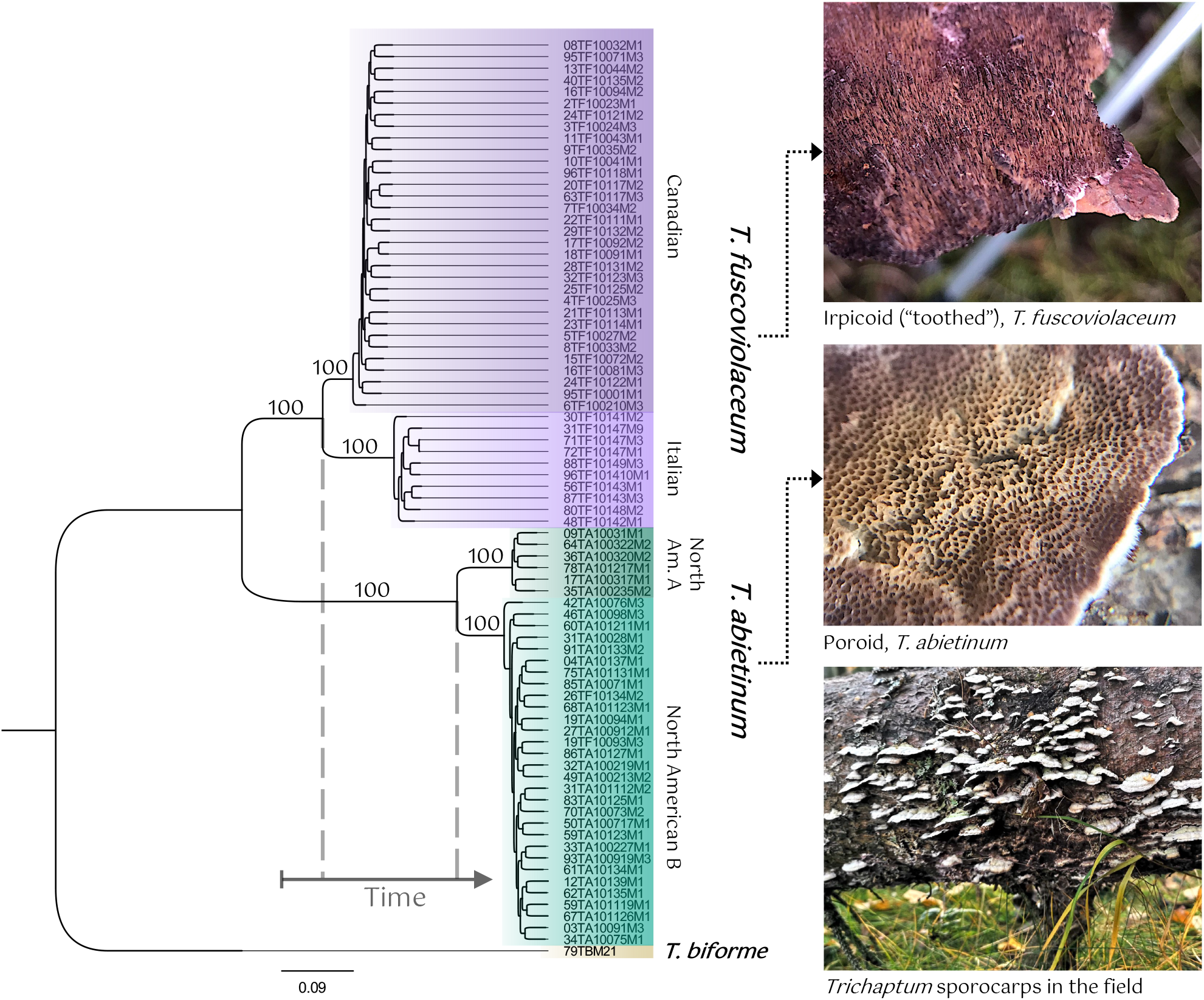
Clear population structure in the phylogenetic tree analysis. The analysis is based on a single nucleotide polymorphism (SNP) dataset of 3 118 957 SNPs. The tree is constructed using *IQ-TREE 2* (Minh et al. 2020) with the model GTR+ASC. The numbers on the branches represent bootstrap branch support. Populations of *Trichaptum fuscoviolaceum* (TF) are coloured in shades of purple and populations of *T. abietinum* (TA) are coloured in shades of green. The outgroup, *T. biforme*, is coloured in brown. The scale bar on the bottom is the number of substitutions per site. The time axis illustrates relative split of the TF and TA populations (see Figure S6). The shade of purple of the hymenium (spore producing layer) can vary. The two TF individuals in the North American B population are confirmed as North American B TA individuals after genomic analyses (wrongly assigned in the field). Photographs of hymenia by Inger Skrede and photograph of sporocarps by Malin Stapnes Dahl.

The PCA also indicated clear groupings of species and populations of *T. fuscoviolaceum* and *T. abietinum*, with PC1 and PC2 explaining 57.5% and 12.2% of the observed variation, respectively (Figure 3). The Italian and Canadian *T. fuscoviolaceum* populations were closer to each other than either was to the North American A and the North American B *T. abietinum* population along PC1. PC2 positioned the Italian *T. fuscoviolaceum* population and the North American A *T. abietinum* population at opposite ends of the axis, while the Canadian *T. fuscoviolaceum* and the North American B *T. abietinum* population were placed closer in the middle of the axis.

**Figure 3.**
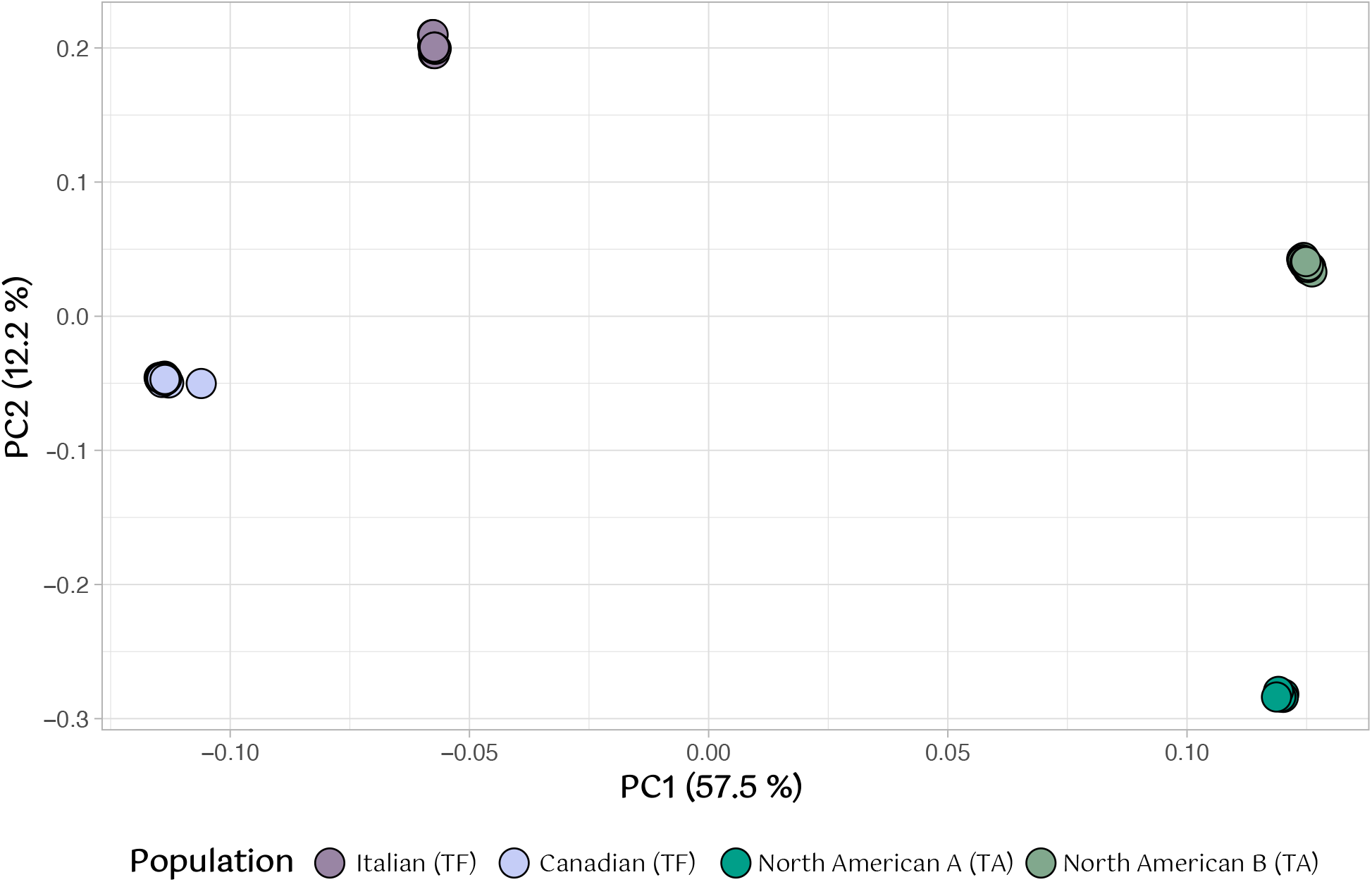
Clear groupings according to species and populations in the principal component analysis (PCA). The PCA is based on a single nucleotide polymorphism (SNP) dataset of 2 040 885 SNPs linkage pruned to 56 046 SNPs. The x and y-axes represent PC1 and PC2, respectively, with percentage of variance explained in parentheses. Points are individuals colored by population as indicated in the legend. TF = *Trichaptum fuscoviolaceum* and TA = *T. abietinum*. The figure is made in *R v4.0.2* using the packages *ggplot2* (Wickham 2016) and *wesanderson* (Ram and Wickham 2018).

The clear distinction spotted in the PCA was corroborated by the fixation index (*F*_ST_), which showed a high degree of divergence both between populations of different species and between populations of same species. The *F*_ST_ means across the genome for between species comparisons were ranging from 0.6 – 0.8. The within-species comparisons showed higher differentiation between the two *T. fuscoviolaceum* populations than between the *T. abietinum* populations (mean *F*_ST_ between *T. fuscoviolaceum* populations was 0.46, while mean *F*_ST_ between *T. abietinum* populations was 0.33).

The absolute between populations divergence (*d*_XY_) echoed the patterns of the PCA and *F*_ST_ scan, with generally high divergence both between populations of different species and between populations within species. The mean *d*_XY_ values between populations of different species were about 0.4, while the mean values between populations of same species were slightly less than 0.2 for both comparisons.

The within population variation calculated by the nucleotide diversity, π, had a mean value of about 0.05 for all populations.

### Introgression analyses indicate a complex evolutionary history

The divergence analyses suggested that the species had diverged for a long time. Thus, we wanted to explore signs of ancestral introgression not revealed by assessing current interbreeding with crossing experiments. The *D* statistic, used to detect signs of introgression across the genome, gave significant *D* values (|z-score| > 3) between the Italian *T. fuscoviolaceum* and both the *T. abietinum* populations (Table 1), indicating introgression between the Italian *T. fuscoviolaceum* and the *T. abietinum* populations. The test of introgression between the *T. abietinum* populations and either of the *T. fuscoviolaceum* populations did not reveal significant positive or negative *D*-values. There was also a larger discrepancy between ABBA and BABA sites in the significant topologies (Table 1, row three and four), than in the nonsignificant topologies (Table 1, row one and two).

**Table 1.** The *D* statistic indicates introgression. The analysis is performed on a single nucleotide polymorphism (SNP) dataset of 3 118 957 SNPs. The *D* statistic is based on a phylogenetic tree hypothesis of (((W, X), Y), Z) and tests introgression between Y and X (negative D) and Y and W (positive D). Z is the outgroup. The table includes the *D* value (*D*), standard error (std error), significance of the *D* values (z-score; an absolute z-score larger than 3 is considered significant), the number of SNPs shared between Y and W (BABA), the number of SNPs shared between Y and X (ABBA), and the number of SNPs used for the comparison (n SNPs). Significant introgression between populations is highlighted in bold.

| ***D* statistics** |  |  |  |  |  |  |  |  |  |
| --- | --- | --- | --- | --- | --- | --- | --- | --- | --- |
| **W** | **X** | **Y** | **Z** | ***D*** | **std error** | **z-score** | **BABA** | **ABBA** | **n SNPs** |
| NAmB TA | NamA TA | Can TF | TB | -0.0041 | 0.004744 | -0.868 | 9142 | 9217 | 662894 |
| NamA TA | NamB TA | It TF | TB | 0.0060 | 0.005929 | 1.018 | 9706 | 9590 | 662712 |
| Can TF | **It TF** | **NamA TA** | TB | -0.2257 | 0.008558 | -26.376 | 8072 | 12776 | 662712 |
| **It TF** | Can TF | **NamB TA** | TB | 0.2287 | 0.009069 | 25.221 | 12810 | 8041 | 664963 |

Can TF = Canadian *Trichaptum fuscoviolaceum*, It TF = Italian *T. fuscoviolaceum*, NamB TA = North American B *T. abietinum*, NamA TA = North American A *T. abietinum*, TB = *T. bifforme*

The four-population *f* statistic (*f*_4_ ratio), used to test proportion of introgression, resulted in a violation of the statistical model (i.e., negative alpha values; valid values are proportions between 0 and 1) when placing *T. abietinum* and *T. fuscoviolaceum* as sister groups with the Canadian *T. fuscoviolaceum* or the North American B *T. abietinum* at the X position (Table 2). Reversing the positions of the Canadian and Italian *T. fuscoviolaceum* or the two *T. abietinum* populations at X and C resulted in a positive alpha value, which did not violate the model (Table 2). The alpha value indicated about 5.7% shared ancestry between the *T. abietinum* populations and the Italian *T. fuscoviolaceum* population (Table 2, row three and four). The small amount of shared ancestry (0.1 – 0.2%) between the North American A *T. abietinum* and the two *T. fuscoviolaceum* populations did not show a significant z-score (< 3; Table 2, row seven and eight).

**Table 2.** The four-population *f* statistic (*f*4) show further signs of introgression. The analysis is based on a single nucleotide polymorphism (SNP) dataset of 3 118 957 SNPs. The table shows the different configurations tested from a hypothesis of the phylogenetic relationship presented as (((A, B), (X, C)), O), where X is the introgressed population and C its sister population, with the B population as the source of introgression and A as its sister population. O is the outgroup (*Trichaptum biforme*). The alpha value indicates proportion of gene flow with standard error (std error) and significance (z-score; considered significant when larger than 3). Negative alpha values are due to violation of the statistical model. Significant introgression between populations is highlighted in bold.

| *F<sub>4</sub>* ratio |  |  |  |  |  |  |  |
| --- | --- | --- | --- | --- | --- | --- | --- |
| A | B | X | C | O | alpha | std error | z-score |
| NamA TA | NamB TA | Can TF | It TF | *T. biforme* | -0.061072 | 0.003619 | -16.878 |
| NamB TA | NamA TA | Can TF | It TF | *T. biforme* | -0.060432 | 0.003553 | -17.010 |
| NamB TA | **NamA TA** | **It TF** | Can TF | *T. biforme* | 0.057001 | 0.003160 | 18.038 |
| NamA TA | **NamB TA** | **It TF** | Can TF | *T. biforme* | 0.057569 | 0.003215 | 17.906 |
| It TF | Can TF | NamB TA | NamA TA | *T. biforme* | -0.002591 | 0.002536 | -1.022 |
| Can TF | It TF | NamB TA | NamA TA | *T. biforme* | -0.001535 | 0.001703 | -0.902 |
| Can TF | It TF | NamA TA | NamB TA | *T. biforme* | 0.001536 | 0.001697 | 0.905 |
| It TF | Can TF | NamA TA | NamB TA | *T. biforme* | 0.002592 | 0.002522 | 1.028 |

NamA = North American A, NamB = North American B, Can = Canadian, It = Italian, TF = *T. fuscoviolaceum*, TA = *T. abietinum*

Further investigation of introgression with the three-population outgroup *f* statistic (*f*_3_), which estimates shared genetic drift (or branch length), revealed that the *T. abietinum* populations split later (share more genetic drift) than the *T. fuscoviolaceum* populations (Figure S6; Figure 2). As with the *f*_4_ ratio analysis, the Italian *T. fuscoviolaceum* population exhibited slightly more shared genetic drift with the *T. abietinum* populations than the Canadian *T. fuscoviolaceum* population. Nevertheless, the difference between the two *T. fuscoviolaceum* populations was miniscule. The *f*_3_ analysis indicated a phylogenetic topology where the *T. fuscoviolaceum* populations diverged earlier than the *T. abietinum* populations. A reasonable next step was therefore to test introgression between the *T. abietinum* populations and each of the *T. fuscoviolaceum* populations in subsequent sliding window introgression analyses (i.e., a (((North American B *T. abietinum*, North American A *T. abietinum*), *T. fuscoviolaceum* population), *T. biforme*) phylogenetic topology).

The network analysis conducted in *TreeMix*, based on the model with the most optimal number of edges, supported introgression from the Italian *T. fuscoviolaceum* into the North American B *T. abietinum* population (Figure 4). From the residual plot (Figure S8, 1 edge), it was clear that a large proportion of the residuals were not accounted for between the Italian *T. fuscoviolaceum* and the North American B *T. abietinum*. When testing which number of edges was the most optimal, the model with 1 migration edge got the best support (Figure S9).

**Figure 4.**
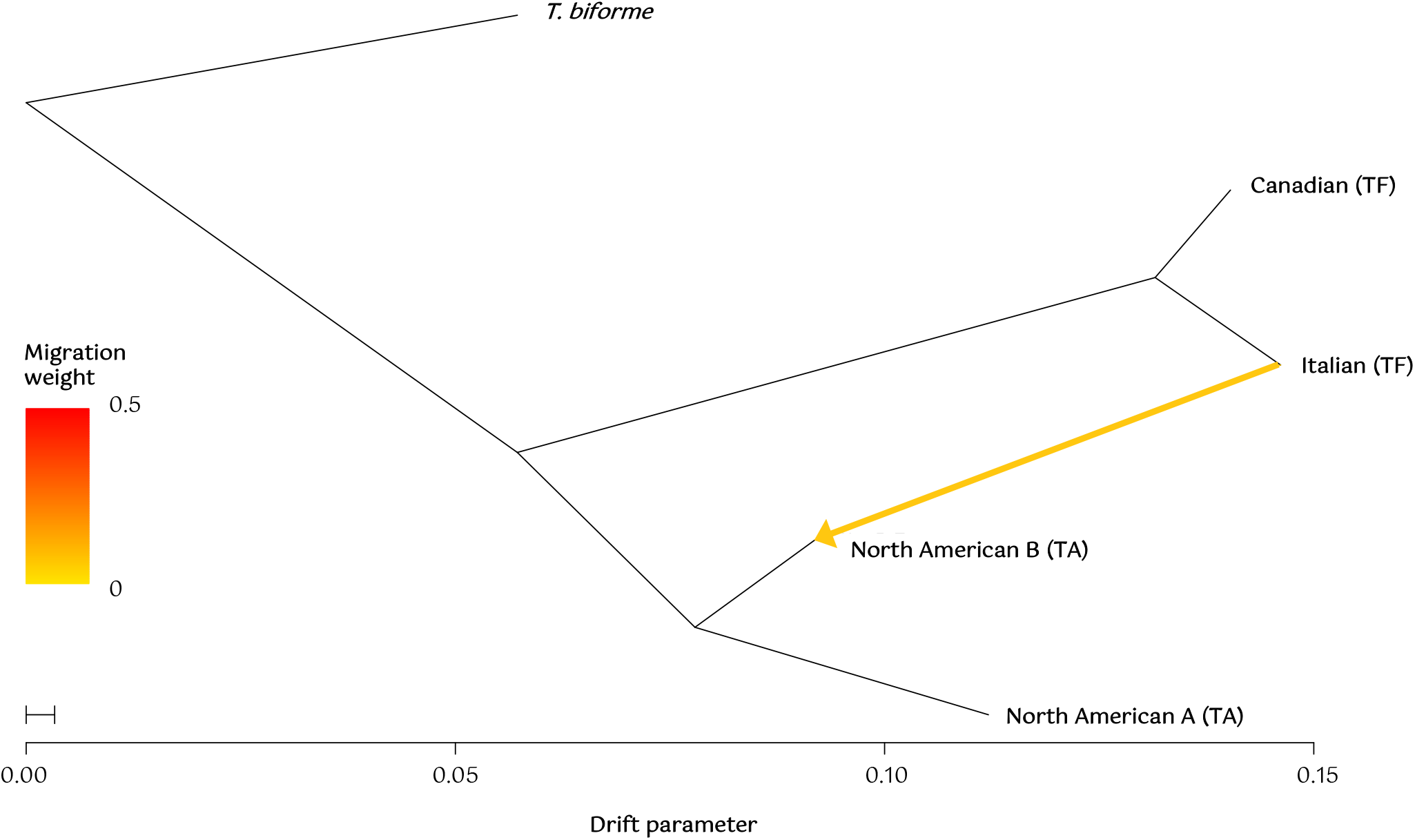
Introgression from the Italian *Trichaptum fuscoviolaceum* into the North American B *T. abietinum*. The plot is based on results from the *TreeMix* (Fitak 2021) analysis with a block size (-k) of 700 and 1 migration edge (-m 1). The yellow arrow shows the direction of migration (introgression). The bar on the left depicts the migration weight (proportion of admixture). The bottom scale bar shows the drift parameter (amount of genetic drift along each population; Wang et al. 2016). TF = *T. fuscoviolaceum* and TA = *T. abietinum*. The outgroup is *T. biforme*.

The sliding window proportion of introgression (*f*_dM_) calculated across the genome, which was set up based on the results from the *f*_3_ analysis, revealed small regions of possible introgression (Figure 5; Figure S10). There were several windows of significant positive *f*_dM_ values (e.g., more shared derived polymorphisms than expected between the *T. fuscoviolaceum* populations and the North American A *T. abietinum* population), which suggests regions of introgressed genes (HMM outliers are marked in Figure 5 and S10, and presented in Table S3). The genes in outlier windows coded for many unknown proteins, but also proteins similar to those found in common model organisms such as *Saccharomyces* spp. and *Arabidopsis thaliana*. The genes with similarity to other organisms are annotated to many different functions (i.e., there are genes involved in oxidoreductases, hydrolases, and transport, among others; The UniProt Consortium 2021). The GO enrichment analysis found some of the HMM outlier genes to be involved in metabolic processes and copper ion transport, to name a few (Table S4). However, after running FDR analysis on the raw p-values, none of the enrichment terms were significant.

**Figure 5.**
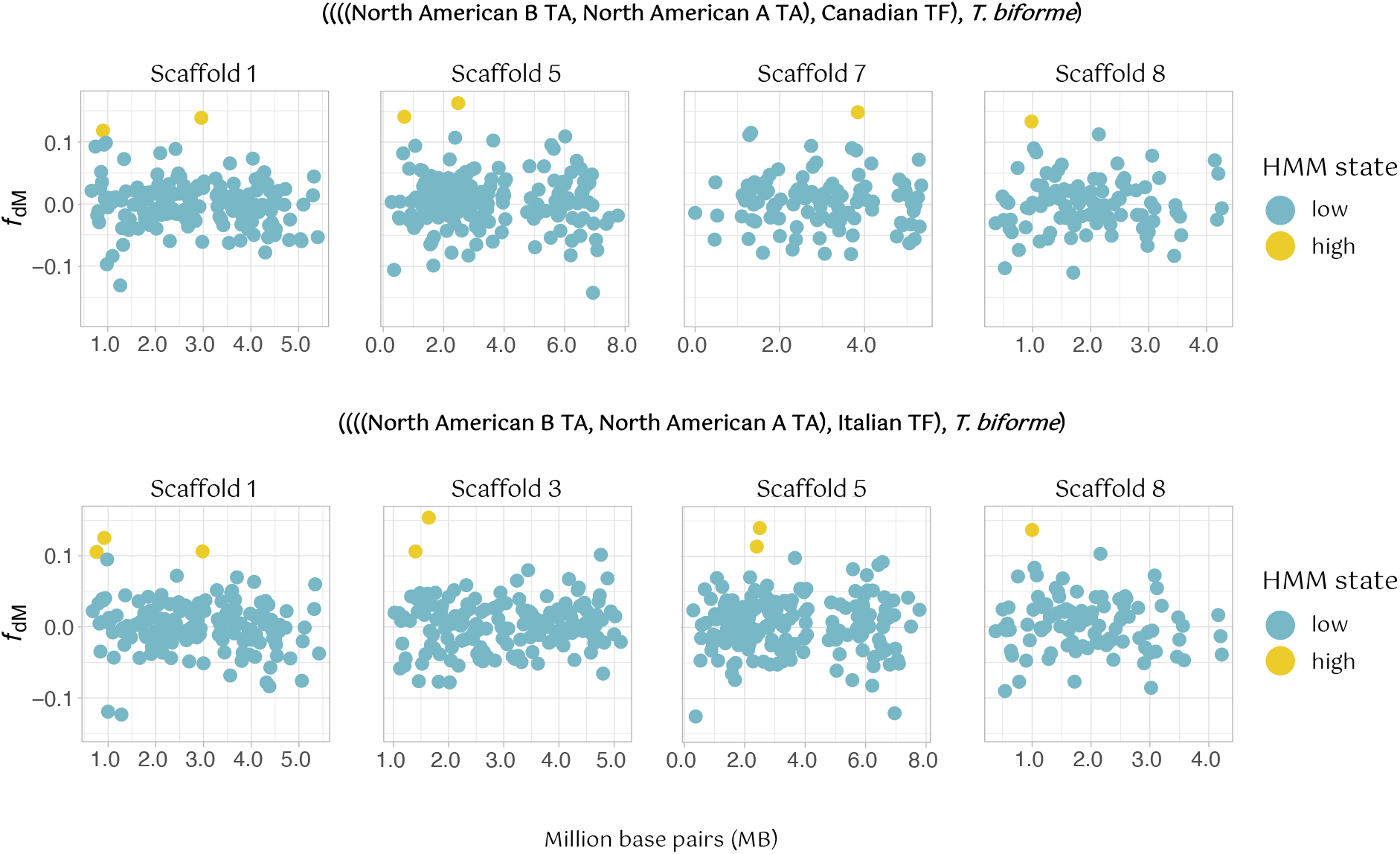
Signs of scattered introgression throughout the genome. A proportion of introgression (*f*_dM_) sliding window analysis based on a single nucleotide polymorphism (SNP) dataset of 3 118 957 SNPs, where windows with at least 100 SNPs are included. Only four of the scaffolds are depicted here for each of the analyses. The remaining scaffolds are shown in Figure S10. The main headers depict the phylogenetic hypothesis, ((((P1, P2), P3), O), where P1, P2 and P3 are populations investigated for introgression and O is the outgroup. A positive value indicates more shared derived polymorphisms than expected between P2 and P3, while a negative value indicates the same for P1 and P3. Each point is the *f*_dM_ value of a window (window size = 20 000 base pairs). Y-axes show the *f*_dM_ value and x-axes represent million base pair (Mb) position of the windows on the scaffolds. The legend shows the Hidden Markov-model (HMM) state of the windows. Blue colored points (low) indicate insignificant amount of introgression, while yellow-colored points (high) are outlier windows with significant introgression from the HMM analysis. Annotated genes in the outlier windows can be found in Table S3. TA = *Trichaptum abietinum* and TF = *T. fuscoviolaceum*. The figure is made in *R v4.0.2* using the packages *tidyverse* (Wickham et al. 2019) and *wesanderson* (Ram and Wickham, 2018).

### Demographic modelling indicates involvement of a ghost population

Lastly, we conducted demographic modelling to gain insight into divergence times and introgression events. We were also able to include a ghost population (Beerli et al. 2004) to test for introgression from unsampled or extinct populations. The best model, supported by both AIC and likelihood distribution comparison, showed introgression occurring twice; first between an ancestral *T. fuscoviolaceum* population and an ancestral *T. abietinum* population and later between a more recent ghost population related to the Italian *T. fuscoviolaceum* and the North American B *T. abietinum* (Figure 6; Figure S7, Ghost migration 9; Figure S11). These results were partly congruent with the network analysis where introgression was inferred between the Italian *T. fuscoviolaceum* and the North American B *T. abietinum* (Figure 4), but in the network analysis a ghost population could not be included. The best supported model further indicated that the sister species split 524 473 generations ago, while the *T. fuscoviolaceum* populations split 99 101 generations ago and the *T. abietinum* populations 92 260 generations ago. The estimated split of the ghost population from the Italian *T. fuscoviolaceum* populations was 94 536 generations ago (Figure 6; Table S5). The analysis further showed that there were few migrants between the ancestral populations and a little more between the ghost population and the North American B *T. abietinum* population, but mostly from the ghost population into the North American B (migration values; Figure 6).

**Figure 6.**
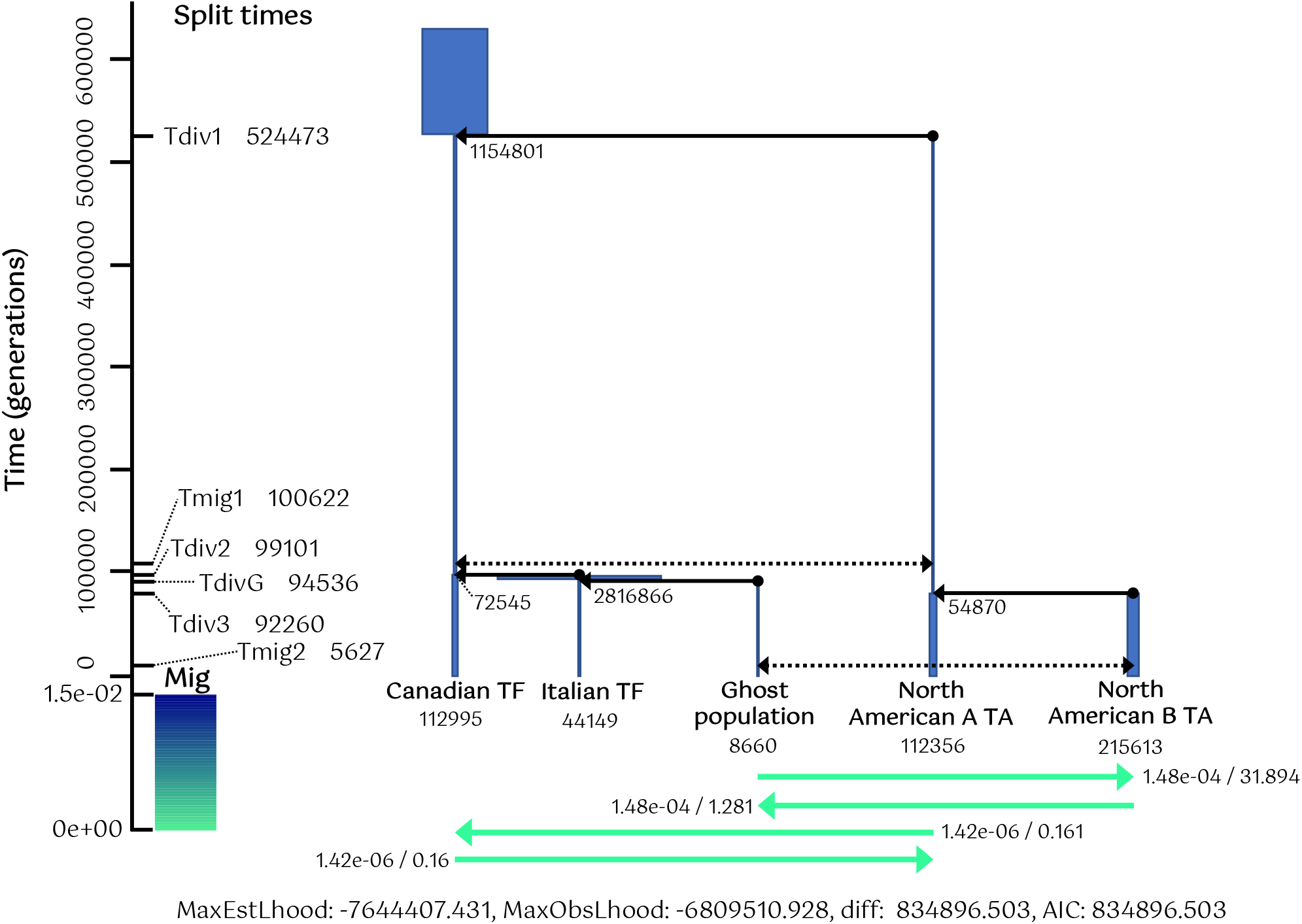
Demographic modelling indicates both ancient introgression and introgression from a recent ghost population. The analysis is performed in *fastsimcoal2* (Excoffier et al. 2021) using a dataset of 3 065 109 SNPs. The figure shows the best supported model. The right side of the axis indicate time in generations, with the lower part (below 0; colour bar) showing proportion of migration. The split times (divergence times) and migration times are presented on the right side of the axis. The blue bars and fully drawn black arrows show the estimated effective population sizes and splits, respectively. Effective population sizes are also written next to the splits and below the current populations. The dotted black arrows are the introgression events. The green arrows display the amount of migration (related to the colour bar) with the proportion of introgression estimate / the calculation to the number from the effective population size (current populations) that migrated. Tdiv = time of divergence, Tmig = time of migration, mig = migration, MaxEstLhood = maximum estimated likelihood, MaxObsLhood = maximum observed likelihood, diff. = difference between MaxEstLhood and MaxObsLhood, TF = *Trichaptum fuscoviolaceum* and TA = *T. abietinum*. Note: All parameters (including migration) are plotted backward in time and in haploid numbers.

## Discussion

### Divergent sister species show signs of introgression

Population genomic and introgression analyses present a window into exploring the dynamics of populations, their genomes and how they evolve. By searching beyond current phylogenies and population structures, intricate evolutionary histories can be revealed. Our study presents several results indicating that high divergence between *T. abietinum* and *T. fuscoviolaceum* does not exclude the possibility of admixture.

Firstly, high divergence values are prevalent throughout the genomes. The large genetic differences between the species can be a result of mechanisms such as reproductive isolation (Nei et al. 1983) and random events over time (i.e., genetic drift; Watterson 1985). The fungi make up an ancient and diverse kingdom that originated over a billion years ago (Berbee et al. 2020), with the oldest fungal-like fossil dating back to 2.4 billion years ago (Bengtson et al. 2017) and an estimate of 2 – 5 million extant species (Li et al. 2021). The divergence of the order Hymenochaetales, which *Trichaptum* belongs to, dates to the Jurassic, about 167 million years ago (Varga et al. 2019). There are 37 accepted species in the genus (Index Fungorum 2021), but no estimates of the age of *Trichaptum*. Seeing the old age of Hymenochaetales and assuming the genus *Trichaptum* is old, time is a likely explanation for the genome wide high divergence between *T. fuscoviolaceum* and *T. abietinum*. The demographic modelling also suggests that the two species split quite some time ago (524 473 generations).

Today, the sister species occur in the same habitat, with similar morphology and ecology, acting as early saprotrophs on newly deceased conifers in the northern hemisphere (Kauserud and Schumacher 2003). As mentioned, the two species can grow on the same host and are sometimes found on exactly the same substrate. However, *T. fuscoviolaceum* is usually found on pine (*Pinus*) and fir (*Abies*; most individuals in this study were collected on balsam fir; *A. balsamea*), while *T. abietinum* is more common on spruce (*Picea*) and larch (*Larix*; Macrae 1967; Peris et al. 2022). Even though habitats overlap, the crossing experiments corroborate previous results (Macrae 1967) in that the sister species do not hybridize in vitro, and the genomic analyses suggest that this does not happen between contemporary populations in the wild either. However, the detection of gene flow between more recent populations in the demographic modelling does imply that mating between species can occur occasionally in nature.

The crossing experiments between *T. fuscoviolaceum* individuals of different populations demonstrate that individuals can still mate successfully even though the divergence analyses exhibit high *F*_ST_ and *d*_XY_ values. The high divergence could be due to geographic separation of the Italian and Canadian population, reducing gene flow between these populations. Compatibility is not observed among all populations of *T. abietinum*, where intersterility is detected between some populations that occur in sympatry (Macrae 1967; Magasi 1976). The genus *Trichaptum* consists of tetrapolar fungi, which means individuals are compatible only when they have different alleles on both of the two mating loci (*MATA* and *MATB*; Fraser et al. 2007; Peris et al. 2022). Previous studies have shown that fungal mating loci are diverse and maintained by balancing selection (May et al. 1999; James et al. 2004), which was recently demonstrated in *Trichaptum* as well (Peris et al. 2022). In *T. abietinum*, additional reproductive barriers other than incompatible mating loci are at play, causing the formation of intersterility groups. However, such barriers can remain incomplete. If reproductive barriers were incomplete during the divergence of *T. abietinum* and *T. fuscoviolaceum*, and they diverged mostly due to genetic drift in geographic isolation, conserved diversity on the mating loci (as observed in Peris et al. 2022) over time can have allowed for introgression by maintaining reproductive compatibility across species. The demographic modelling does suggest that introgression happened quite some time after the split between the species. This again supports allopatric divergence, making it possible for the species to reproduce at a later stage due to the possible lack of reproductive barriers (no reinforcement). Gene flow between divergent species of fungi has been detected before (e.g., Maxwell et al. 2018). This could be facilitated by the flexible developmental biology of some fungi, with the capability of tolerating developmental imprecision and distortion of their genetic makeup and still be able to grow and reproduce (Moore et al. 2011; Stukenbrock 2016).

The specific mechanisms behind how the two species diverged are difficult to untangle based on our results. However, the *D* and *f* statistics, together with the network analysis and demographic modelling, show signs of introgression between *T. abietinum* and *T. fuscoviolaceum*. Based on the *D* statistic, the ancestor of the Italian *T. fuscoviolaceum* population appears to have admixed with the *T. abietinum* populations. This might be somewhat counterintuitive, as it is the Canadian *T. fuscoviolaceum* population that currently occurs in sympatry with the collected *T. abietinum* populations. However, reproductive barriers can be produced between species in sympatry due to reinforcement (Abbott et al. 2013). When an allopatric lineage, such as the Italian *T. fuscoviolaceum* or a closely related ghost population, is encountered, reproductive barriers may not be in place and gene flow can occur. The *f*_4_ ratio test further corroborates these results, indicating that the Italian *T. fuscoviolaceum* shares a larger proportion of the genome with the *T. abietinum* populations than the Canadian *T. fuscoviolaceum*. The violation of the statistical model for some topologies with *T. abietinum* and *T. fuscoviolaceum* populations as sister species in the *f*_4_ ratio test can be due to lack of data from populations not sampled (extinct and extant; i.e., ghost populations; Beerli 2004), suggesting a more complex evolutionary history of *Trichaptum* than the collected data can disclose. This is corroborated by the demographic modelling, where the best model includes introgression between a ghost population related to the Italian *T. fuscoviolaceum* and the North American B *T. abietinum* population. According to the network analysis, introgression has occurred from the Italian *T. fuscoviolaceum* population into the North American B *T. abietinum* population. However, *TreeMix* is not always able to reveal the true introgression scenario when the actual admixed populations are related to the populations used in the analyses (Fitak 2021). Therefore, introgression has not necessarily occurred between these two populations but most likely between the ghost population incorporated in the demographic modelling and the North American B *T. abietinum* population. This is similar to introgression inferred between archaic hominins, such as Denisovans and Neanderthals, and present-day humans (Durvasula and Sankararaman 2019). A wider collection, including more populations across the northern hemisphere (e.g., Asia and throughout Europe and North America), could capture the ghost population (if extant) and help untangle the shared evolutionary history of *T. abietinum* and *T. fuscoviolaceum*.

The *f*_dM_ analysis shows only slightly more significantly introgressed regions between the Italian *T. fuscoviolaceum* population tested against the *T. abietinum* populations than the Canadian *T. fuscoviolaceum* (14 vs. 11). All the introgressed regions occur between the *T. fuscoviolaceum* populations and the North American A population (not including B as in the other analyses). The demographic modelling did detect introgression between an ancestral *T. fuscoviolaceum* population and a *T. abietinum* population leading up to the current North American A population. Since this event is ancient, introgressed genes have had time to spread and become fixed in the genomes of current populations (in this case the North American A population and the *T. fuscoviolaceum* populations), which could explain why significant regions are only detected between the North American A population and the *T. fuscoviolaceum* populations.

Many of the genes are also found in the same regions across the genome for both comparisons. This further suggests that the *f*_dM_ analysis is detecting ancestral and not recent gene flow because the regions are conserved through the population splits (i.e., the introgression happened before the current populations diverged). The small regions of scattered introgression in the *f*_dM_ analysis also imply more ancient introgression. This follows similar patterns with highly divergent genomes and localized regions of introgression as found in analyses of three-spined stickleback species pairs in the Japanese archipelago (Ravinet et al. 2018) and *Heliconius* butterflies in Brazil (Zhang et al. 2016).

### Population histories, introgression and its implications

In this study, we only have four populations of two widespread species, thus we are not covering the full diversity of the species. Still, the analyses are able to detect intricate population histories including both a population that is not sampled and ancestral populations leading up to the current ones (i.e., ghost populations). The chance of being able to sample all populations or have no ghost populations in an evolutionary study system consisting of natural populations is minor. It is therefore promising that we are able to extract interesting results based on a relatively small sample. The method is also useful for organismal groups such as fungi, where ancient genomes cannot be retrieved due to poor fossilization (Berbee et al. 2020; ancestral populations have for example been detected through genomic sequencing of subfossil in a study of the giant panda; Sheng et al. 2019). Including ghost populations in modelling can improve the estimate of migration rates (Beerli et al. 2004; Slatkin 2004). To increase our understanding of how introgression and gene flow affects the speciation continuum, it requires that researchers account for scenarios such as extinct lineages and ghost populations when performing model testing.

The signs of introgression observed in the oldest migration in the demographic modelling and in the *D* and *f* statistics are likely a case of introgression from extinct lineages. Ancient introgression has previously been detected from extinct cave bears in the genomes of brown bears (*Ursus arctus*; Barlow et al. 2018), through phenotype analyses of beak sizes in one of Darwin’s finches (*Geospiza fortis*; Grant and Grant 2021), and in the mitochondrial genome of the intermediate horseshoe bat (*Rhinolophus affinis*; Mao et al. 2012), to name a few. Genes or alleles from extinct lineages can therefore persist in extant species and might impose adaptive benefits (The *Heliconius* Genome Consortium et al. 2013; Racimo et al. 2015). It is difficult to say if this is the case with *T. abietinum* and *T. fuscoviolaceum*, but genes found in HMM outlier windows of the *f*_dM_ analysis may represent putatively adaptive genes with an ancient introgression origin. Many of the genes code for proteins of unknown function, which is common in non-model organisms due to limited research. However, the genes with similarity to other functional annotated genes are involved in several different functions in organisms. For example, oxidoreductases and hydrolases partake in numerous enzymatic reactions and are known to be important for wood decaying fungi to depolymerize the recalcitrant woody substrate (Floudas et al. 2012; Presley and Schilling 2017). Nevertheless, whether any of these genes are involved in adaptive introgression cannot be concluded based on the *f*_dM_ analysis alone. To extrapolate any adaptive implications from the GO enrichment analysis would not be appropriate, based on the lack of significance. It is also possible that the signs of introgression observed in the *f*_dM_ analysis are due to non-adaptive factors. For example, parts of the genome stemming from ancient introgression can persist due to recombination and constraint (e.g., Schumer et al. 2016; 2018). Thus, this question remains inconclusive until further analyses are conducted (e.g., recombination rates along the genome). However, the retention of the introgressed regions in the genome is still interesting and acts as a detection marker for the evolutionary history of these species not revealed by examining compatibility in the current populations.

Genes transferred through introgression can lead to an expansion of a species’ distribution range, as for example seen for habitat and climate adaptation in cypress species (*Cupressus* spp.; Ma et al. 2019). The divergence of many of the taxa in the family Pinaceae, which includes the current host species of *T. fuscoviolaceum* and *T. abietinum*, is dated to the Jurassic (< ∼185 million years ago; Ran et al. 2018), the same period as the divergence of Hymenochaetales (Varga et al. 2019). There are several examples where research on cryptic diversity in fungi has revealed high divergence and old divergence times when species initially were thought to be closely related (summarized in Skrede 2021), which may also be the case for *T. fuscoviolaceum* and *T. abietinum*. Our results do not conclusively show adaptive introgression but based on the large nucleotide discrepancies and most likely old divergence, one could speculate that the introgression from ancestral populations has facilitated adaption to a larger host range of *T. fuscoviolaceum* and *T. abietinum* as conifers diverged and expanded across the northern hemisphere. However, additional research (e.g., protein function analysis) is needed to say anything certain about the implications of introgression between the sister species.

Since introgression can have impacts on the evolutionary trajectory of a species, it is an important mechanism to consider when investigating the evolutionary history of taxa. Introgression is not well examined in fungi or within an experimental system based on natural populations, and historically most introgression studies have been carried out on mammals or plants (Dagilis et al. 2021). Our results indicate that ancient introgression can be detected also among divergent species. Even though the phylogenetic relationship between *T. fuscoviolaceum* and *T. abietinum* is well-defined (Seierstad et al. 2020; Peris et al. 2022), signals of introgression lingering in their genomes suggests that the evolutionary history of these species is more complex than the current phylogenies can reveal.

## Conclusion

Our study corroborates earlier findings, indicating that *T. abietinum* and *T. fuscoviolaceum* do not hybridize in vitro. Our results show that the sister species are highly divergent, exhibiting large genetic differences and are reproductively isolated. Nevertheless, introgression analyses display admixture, with small regions of introgression occurring throughout the genomes. These signs point to cases of both ancient and recent introgression between ancestral and current populations of *T. abietinum* and *T. fuscoviolaceum*, including a ghost population of a non-sampled or extinct population. Regardless of a well-resolved phylogeny, the evolutionary history of these species is intricate, including transfer of genes across lineages with unknown implications. This study builds on a small collection of studies detecting introgression between highly divergent species, expanding our knowledge on speciation and the permeability of reproductive barriers. The study also presents a novel system including natural populations and in vitro experiments, which is much needed for understanding the speciation continuum. The ceaselessness of speciation will naturally leave traces of historical events in the genomes of extant organisms. Accounting for these events when investigating speciation and adaptation can give insight into how evolution proceeds and shapes the diversity we observe today, as well as how populations are affected in the future. It will be interesting to use this fungal experimental system applying other approaches, including protein function analysis, to link introgression to historical events (e.g., host shifts) and increase insight into the mechanisms governing divergence and adaptation.

