## Supporting information for "Introgression between highly divergent fungal sister species"

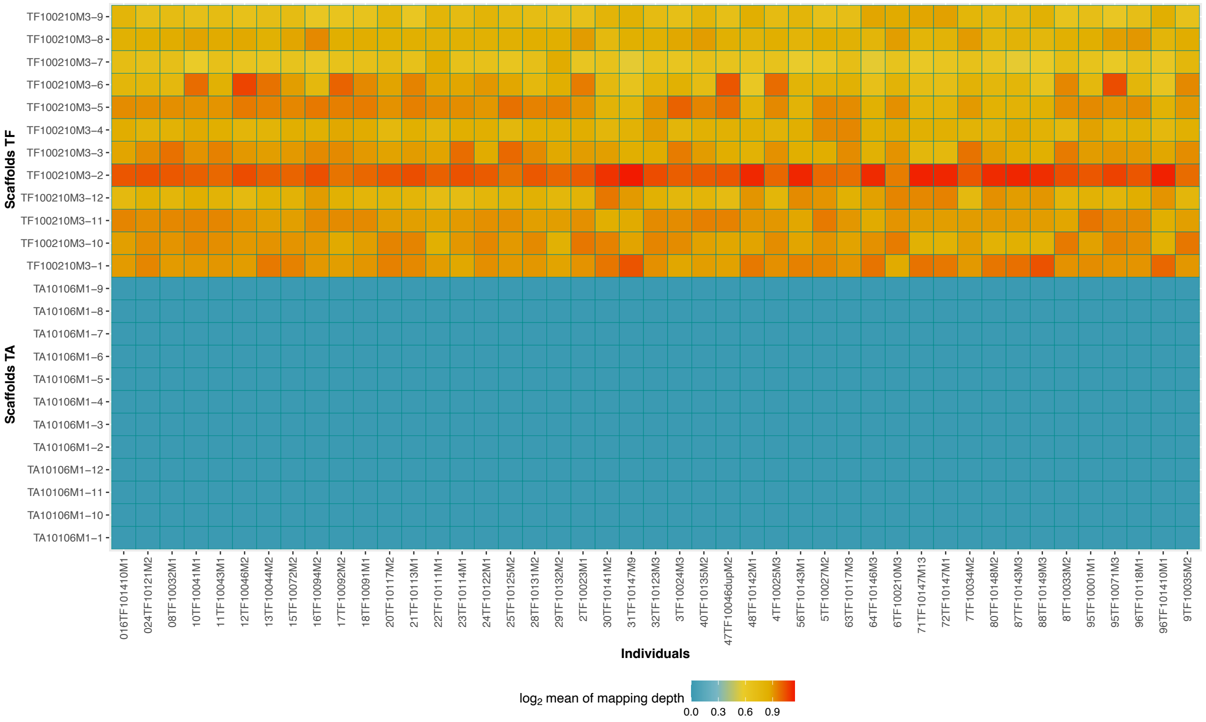


**Figure S1**. **The *sppIDer* analysis did not detect hybrid individuals.** On the y-axis are scaffolds (chromosomes) of the *Trichaptum fuscoviolaceum* (TF) and *T. abietinum* (TA) combined reference genome. The x-axis shows *T. fuscoviolaceum* individuals mapped to the combined reference genome. The legend on the bottom shows a colour gradient for the $\log_2$ mean of the mapping depth. Cooler colours indicate poorer mapping, while warmer colours indicate better mapping. The figure is made in *R v4.0.2* (R Core Team) using the packages *ggplot2* (Wickham, 2016), *wesanderson* (Ram and Wickham, 2018), *viridis* (Garnier, 2018), *readtext* (Benoit and Obeng, 2020), *data.table* (Dowle and Srinivasan, 2020) and *hrbrthemes* (Rudis, 2020).

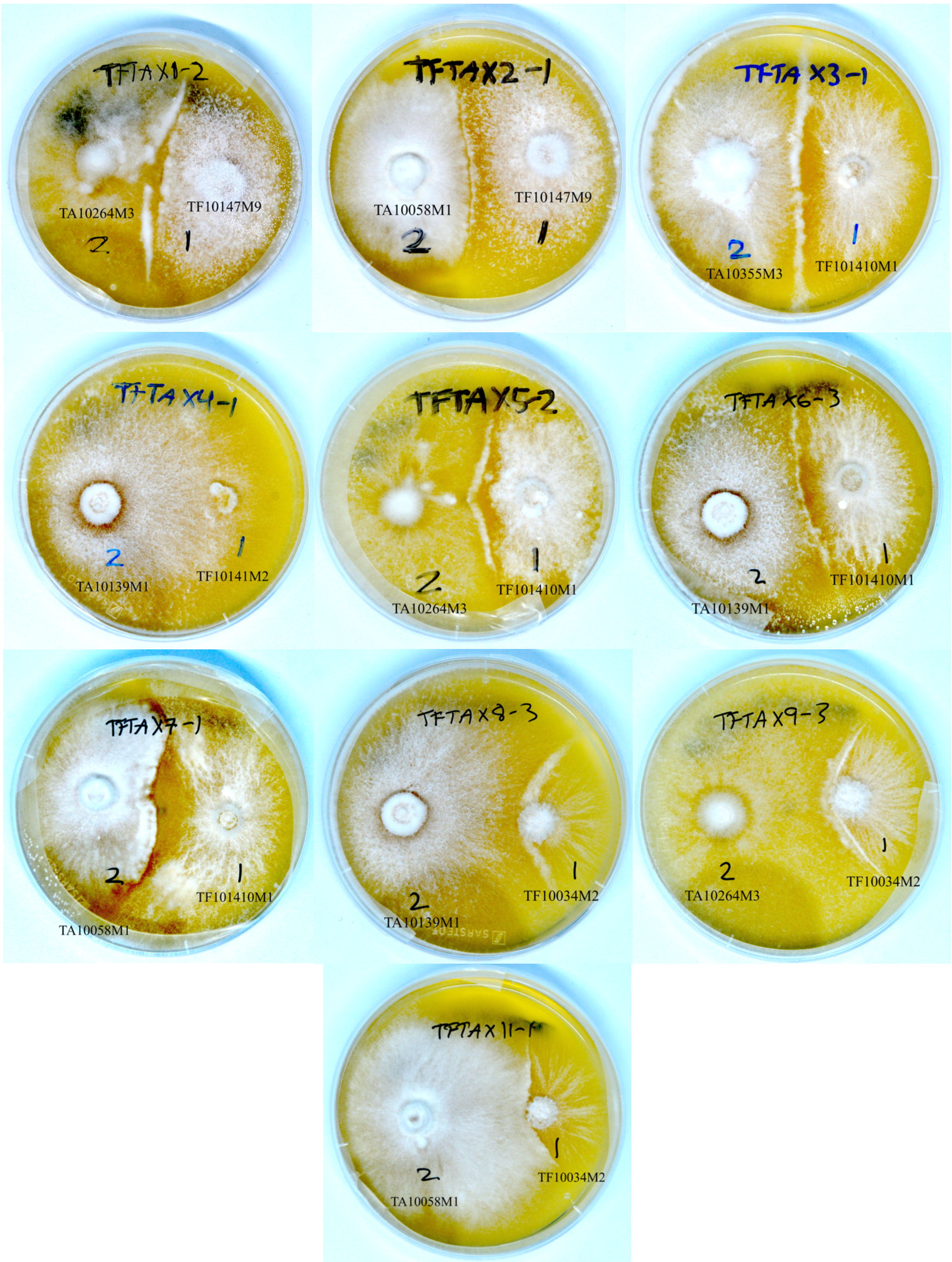


**Figure S2. No successful crossings between *Trichaptum abietinum* and *T. fuscoviolaceum*.** Photographs of cultures from the crossing experiments (one of the three replicates for each cross). None of the crossings are successful. The cross name is indicated at the top (number after the dashed line indicates replicate number), and the individuals are noted on the bottom. TA = *T. abietinum* and TF = *T. fuscoviolaceum*. Photographs were taken with a Nikon D600 Digital Camera (Tokyo, Japan).

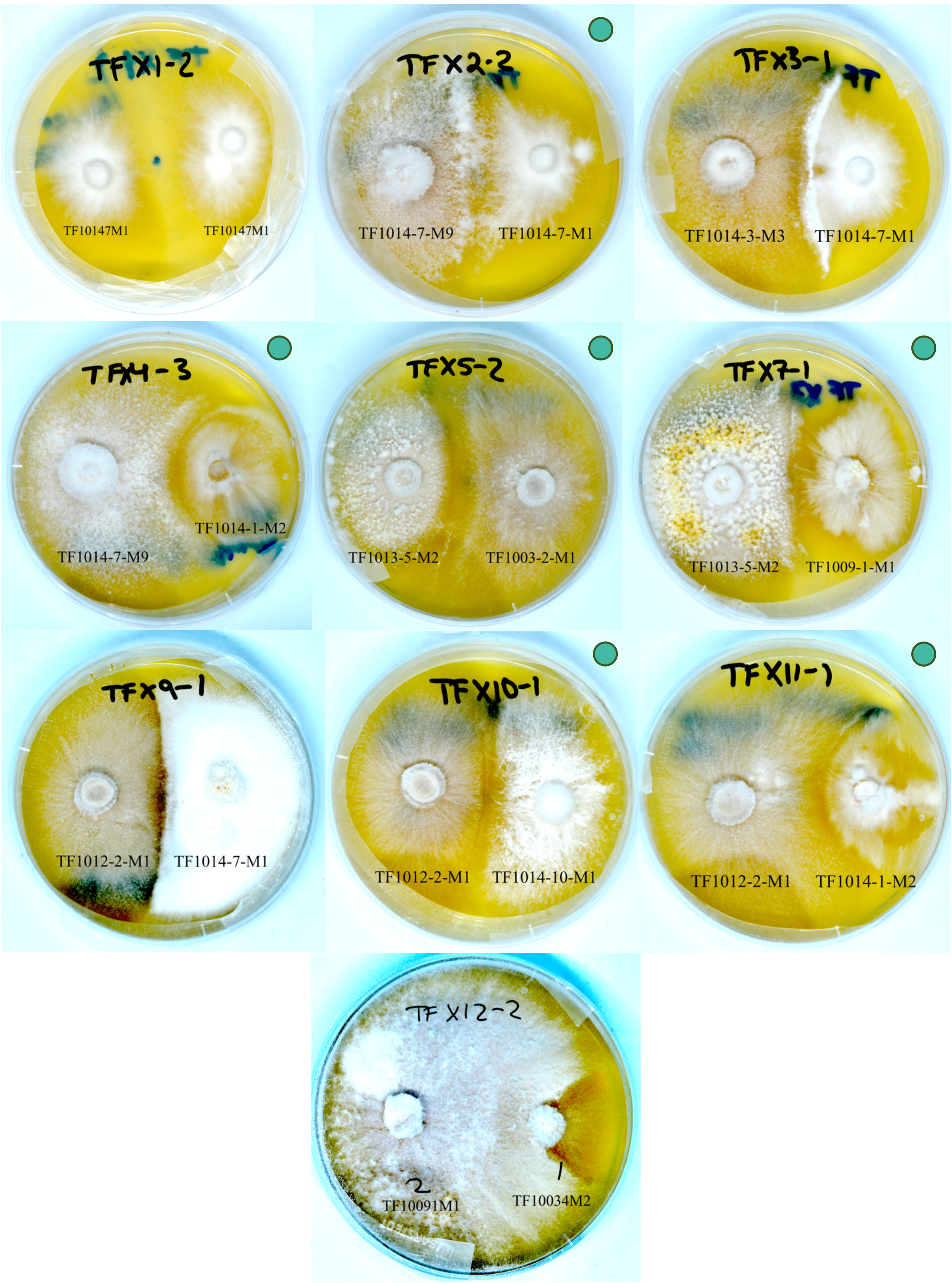


**Figure S3. Crossings between individuals of *T. fuscoviolaceum* mated as expected.** Photographs of cultures from the crossing experiments (one of the three replicates for each cross). Crossings between individuals were as expected (see Table S2). Successful crossings are marked with a green circle. The cross name is indicated at the top (number after the dashed line indicates replicate number), and the individuals are noted on the bottom. TF = *T. fuscoviolaceum*. Photographs were taken with a Nikon D600 Digital Camera (Tokyo, Japan).

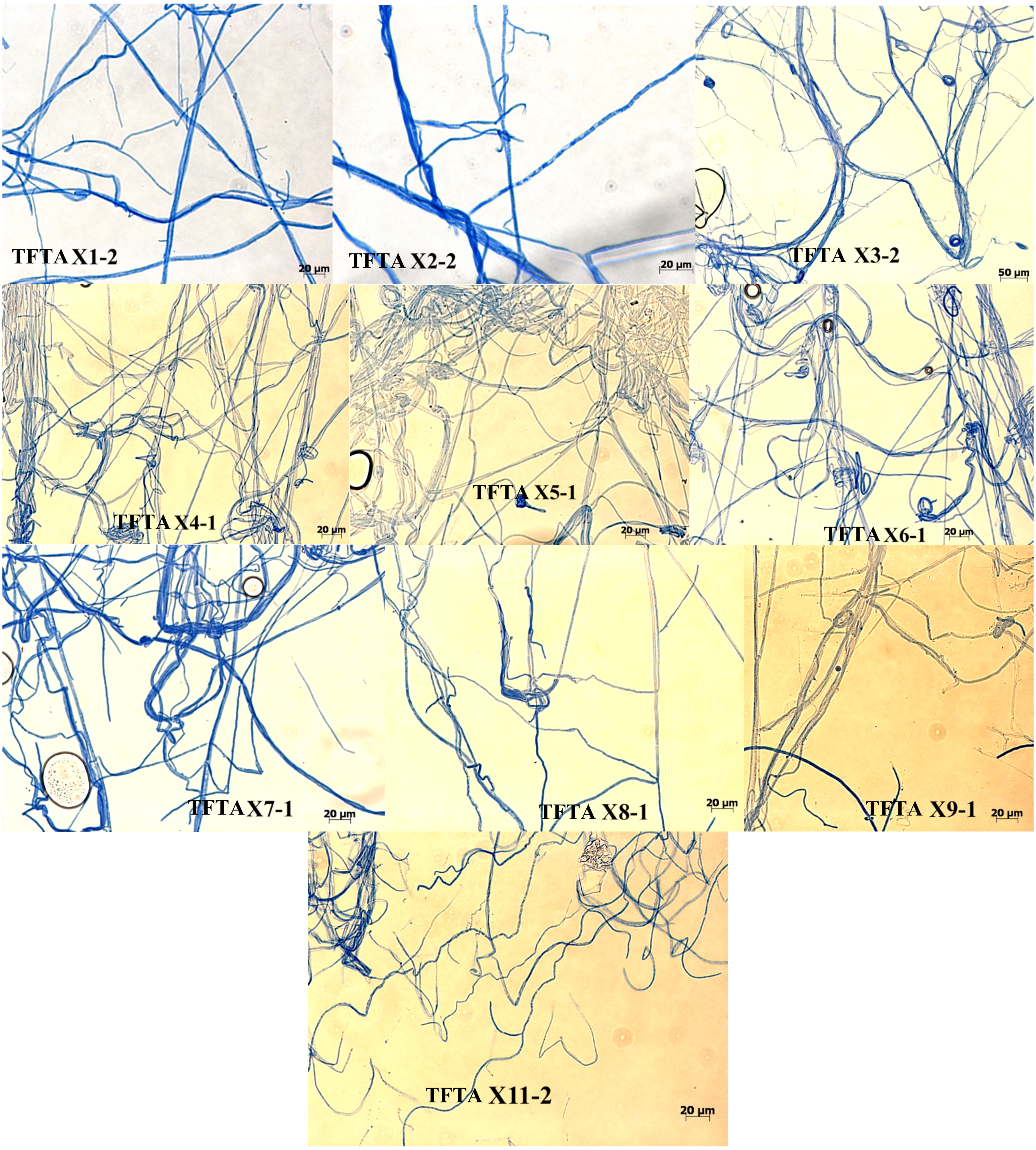


**Figure S4. No clamp connections in the crossings between *Trichaptum abietinum* and *T. fuscoviolaceum*.** Microscope photographs of the crossing experiments. Cross names are indicated at the bottom of the pictures and the number after the dashed lines indicate replicate number. A scale bar is positioned at the bottom right of every picture. TA = *T. abietinum* and TF = *T. fuscoviolaceum*. Photographs were taken using Zeiss Axioplan 2 imaging light microscope (Güttingen, Germany) with Zeiss AxioCam HRc (Güttingen, Germany).

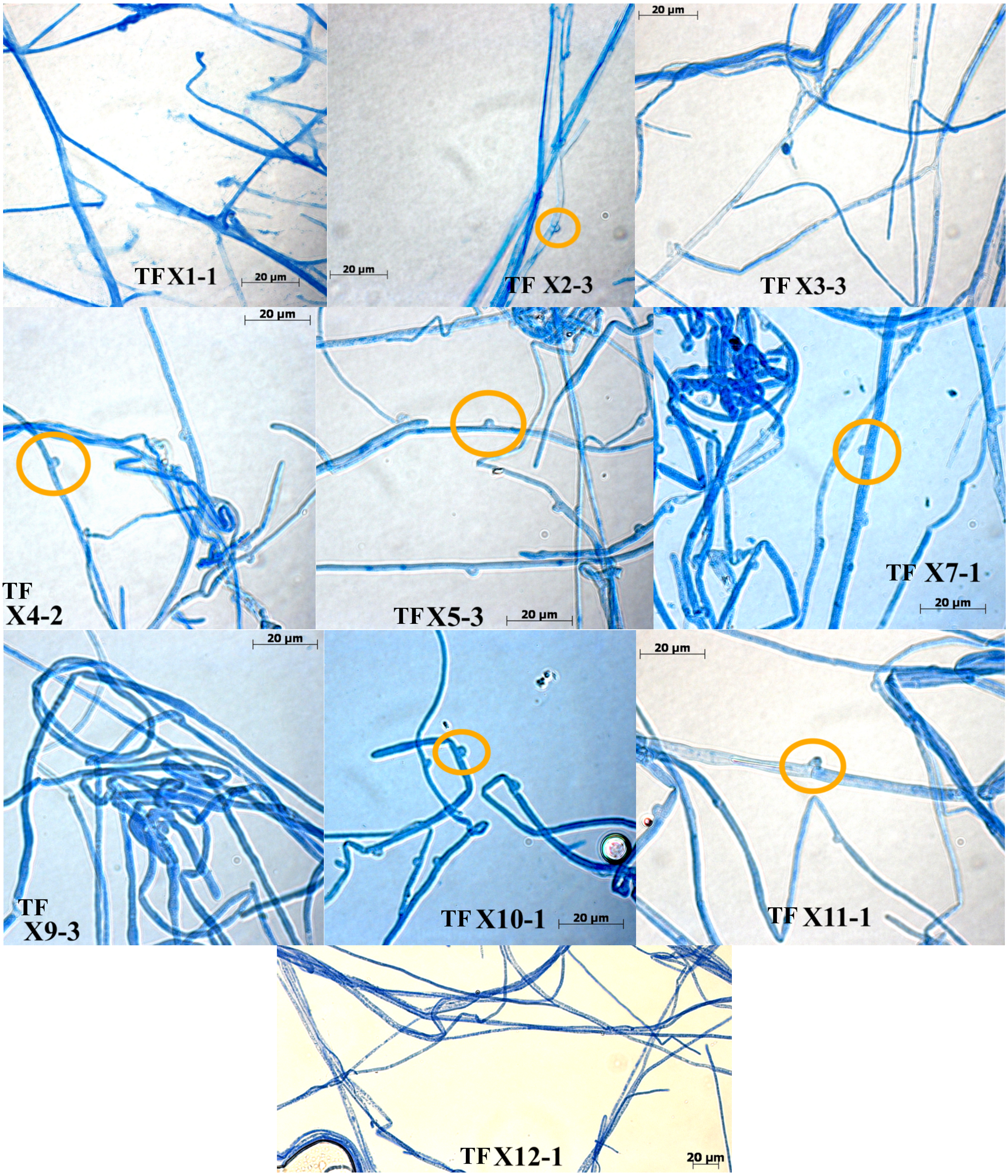


**Figure S5. Clamp connections between successful matings as expected in the crossings between individuals of *T. fuscoviolaceum*.** Microscope photographs of the crossing experiments. Clamp connections are marked with an orange circle. Cross names are indicated at the bottom of the pictures and the number after the dashed lines indicate replicate number. A scale bar is positioned at the bottom right of every picture. TF = *T. fuscoviolaceum*. Photographs were taken using Zeiss Axioplan 2 imaging light microscope (Güttingen, Germany) with Zeiss AxioCam HRc (Güttingen, Germany).

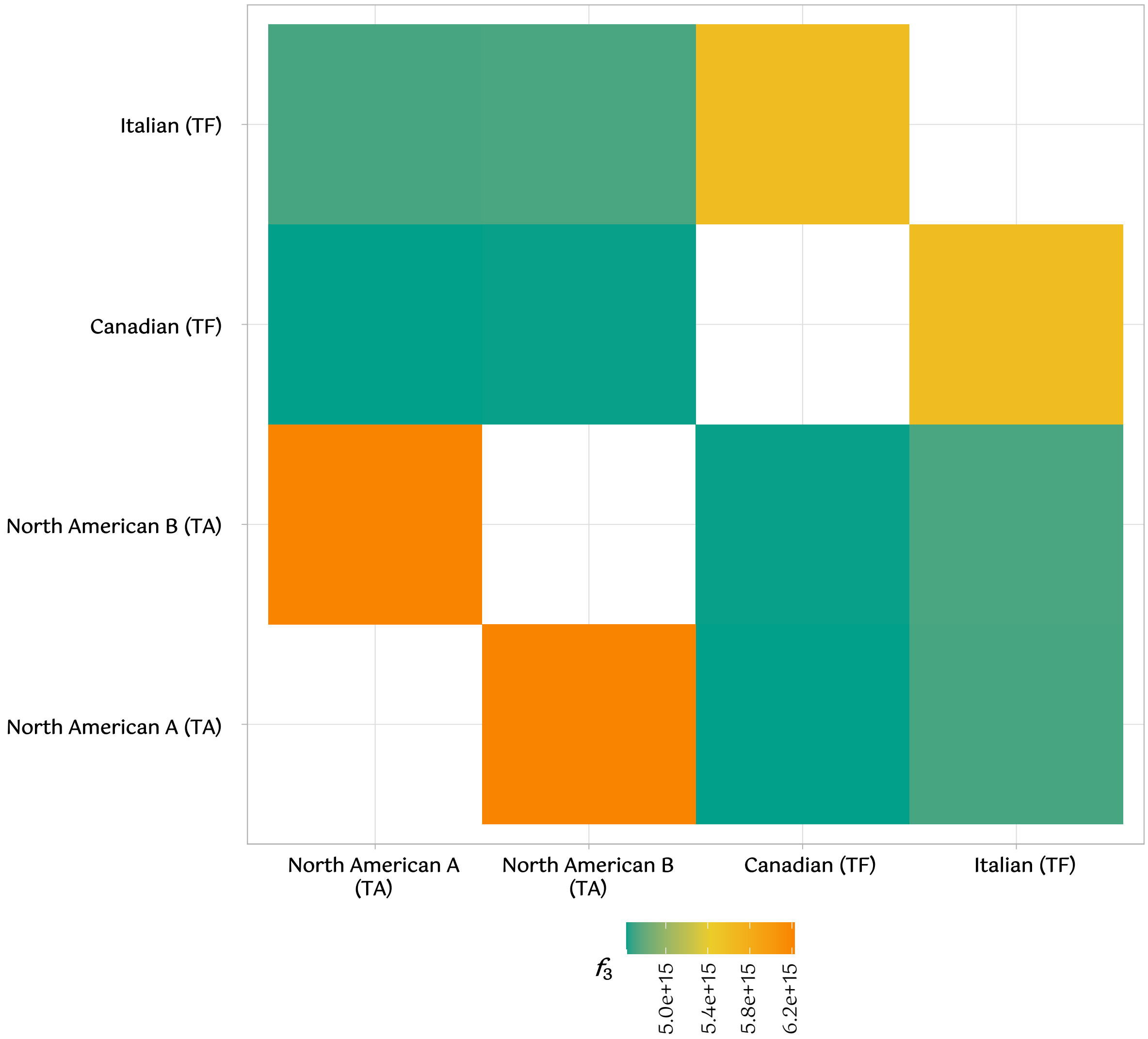


**Figure S6**. ***Trichaptum abietinum* populations split more recently than the *T. fuscoviolaceum* populations in the three-population *f* statistic ($f_3$) with outgroup.** The analysis is based on a single nucleotide polymorphism (SNP) dataset of 3 118 957 SNPs. The figure shows a pairwise comparison of populations colored by amount of shared evolutionary history. The analysis is based upon the phylogenetic hypothesis ((A, B), C)), where the branch lengths of A and B are estimated relative to C. All populations have been tested at position A and B, while C is kept constant as the outgroup *Trichaptum biforme*. The color legend at the bottom depicts relative split (amount of shared genetic drift) between the two populations compared. Higher values indicate a later split than lower values. In Figure 2 these results are illustrated by the time arrow and dotted lines showing the relative split of *T. fuscoviolaceum* populations compared to *T. abietinum* populations from their common ancestor. TF = *T. fuscoviolaceum* and TA = *T. abietinum*. The figure is made in *R v4.0.2* using the packages *admixr* (Petr et al. 2019), *tidyverse* (Wickham et al. 2019) and *wesanderson* (Ram and Wickham 2018).

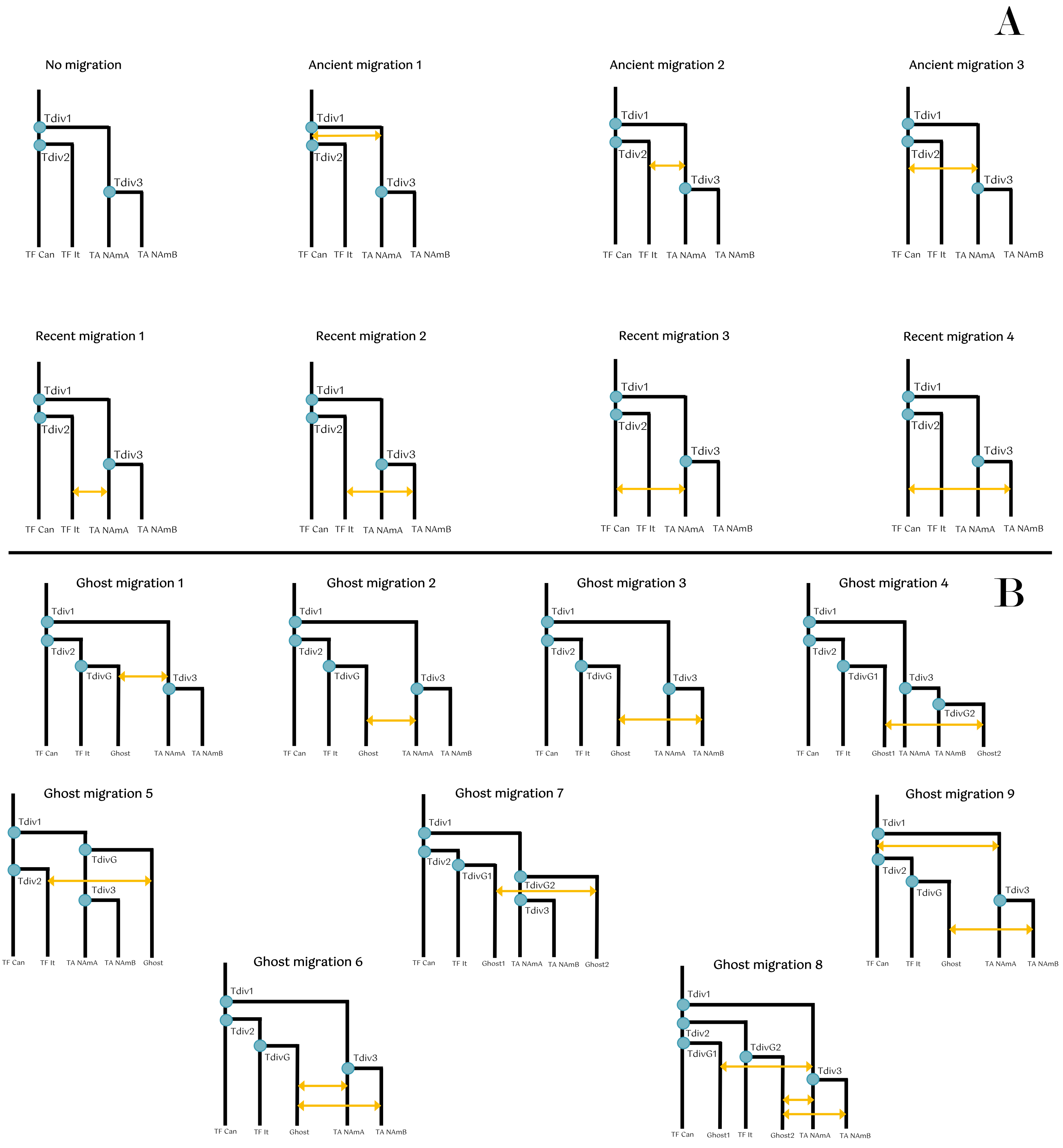


**Figure S7. Models used to test divergence and introgression (migration) times.** Illustrations of how the different models tested were set up in *fastsimcoal2* (Excoffier et al. 2021). Blue dots are the divergence times being estimated and the yellow arrows are the migrations times being estimated. (A) Models without ghost populations. (B) Models with ghost populations. Tdiv = time of divergence, TF = *Trichaptum fuscoviolaceum*, TA = *T. abietinum*, Can = Canadian, It = Italian, NAmA = North American A and NAmB = North American B.

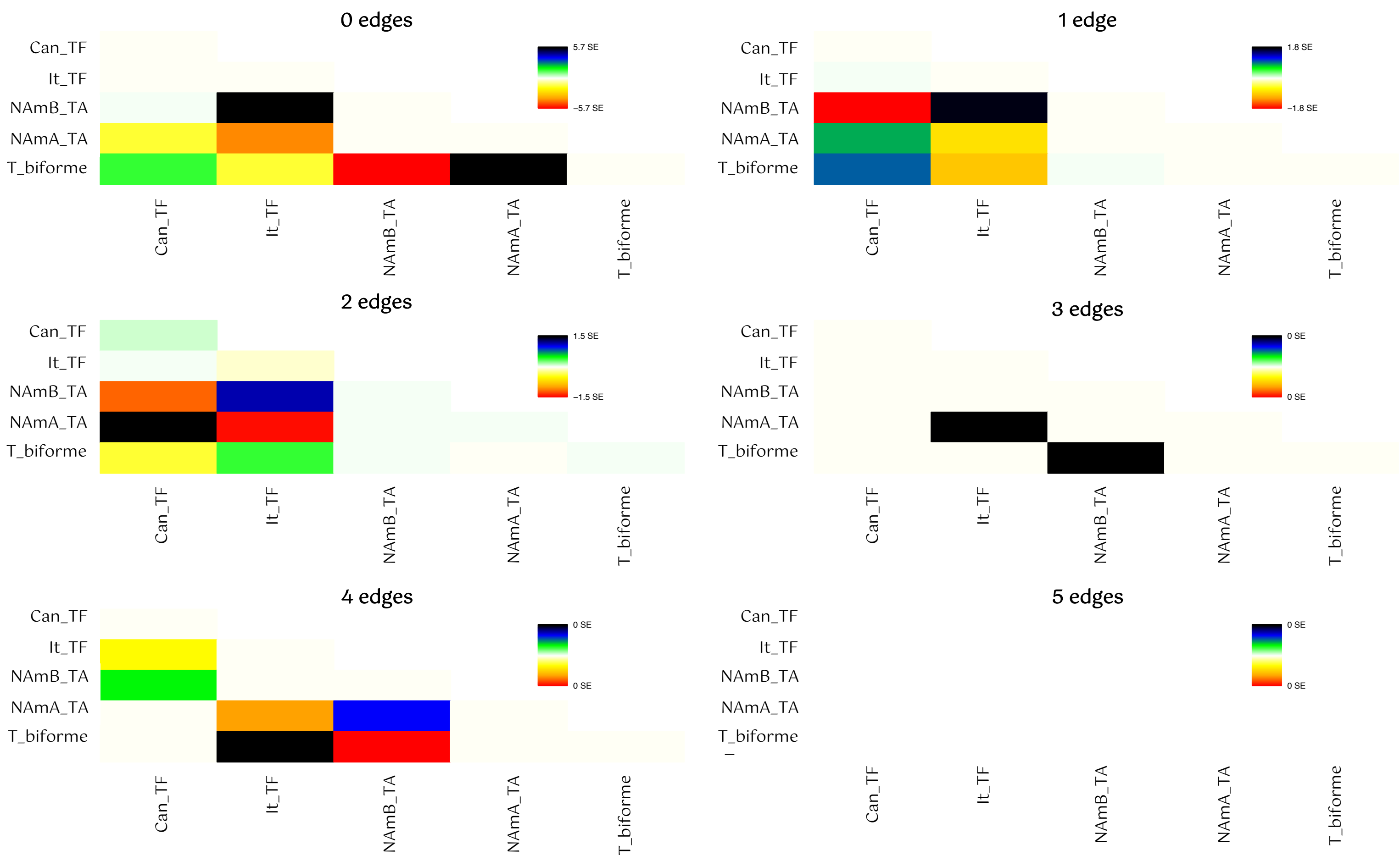


**Figure S8. Residual plots from the *TreeMix* analysis.** Residuals plotted for different edges based on the *TreeMix* (Fitak 2021) analysis using a block size of 700. The colour bar on the right shows the standard error (SE). A higher SE (e.g., black square) indicates that a large portion of the residuals are not accounted for in the model. TF = *Trichaptum fuscoviolaceum*, TA = *T. abietinum*, Can = Canadian, It = Italian, NAmA = North American A and NAmB = North American B. *T. biforme* is the outgroup.

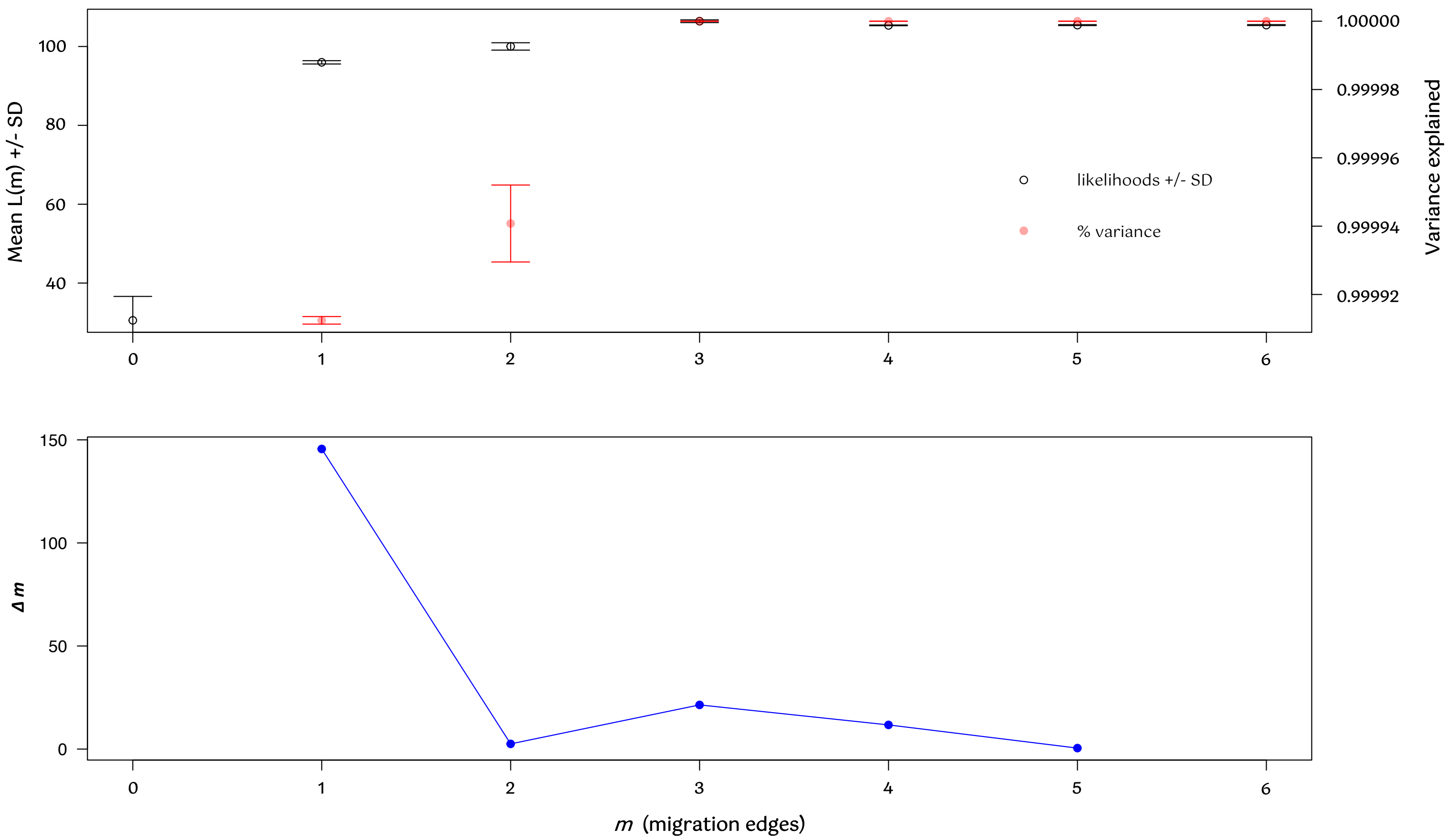


**Figure S9. The *optM* (Fitak 2021) analysis indicates that the *TreeMix* model with 1 edge is the most optimal.** The upper panel shows the mean and standard deviation for the composite likelihood on the left y-axis and the proportion of variance explained on the right y-axis. The bottom panel depicts the second-order rate of change (*Δm*) across values of *m* on the y-axis. The x-axis in both panels indicates the number of migration edges. The peak at edge 1 in the bottom panel is considered to represent the most optimal edge number.

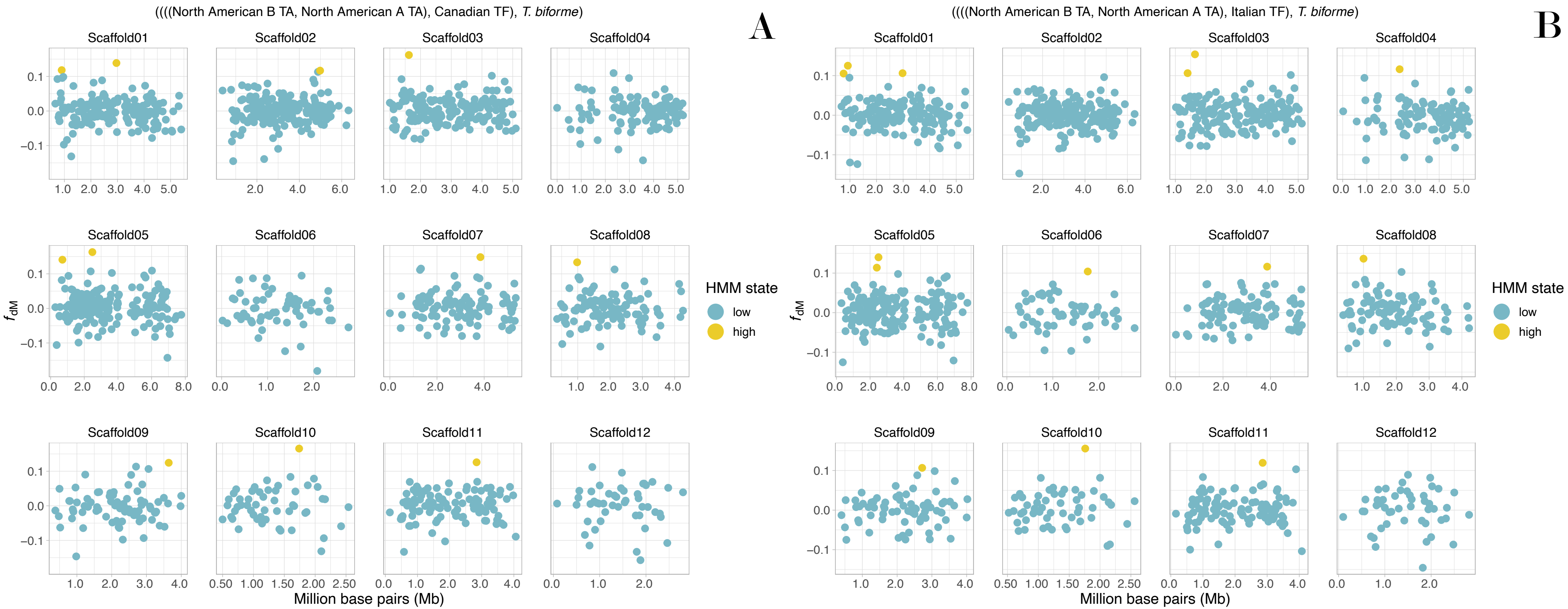


**Figure S10. Signs of scattered introgression throughout the genome**. A proportion of introgression ($f_{dM}$) sliding window analysis based on a single nucleotide polymorphism (SNP) dataset of 3 118 957 SNPs, where windows with at least 100 SNPs are included. The main headers depict the phylogenetic hypothesis, ((((P1, P2), P3), O), where P1, P2 and P3 are populations investigated for introgression and O is the outgroup. A positive value indicates more shared derived polymorphisms than expected between P2 and P3, while a negative value indicates the same for P1 and P3. Each point is the $f_{dM}$ value of a window (window size = 20 000 base pairs). Y-axes show the $f_{dM}$ value and x-axes represent million base pair (Mb) position of the windows on the scaffolds. The legend shows the Hidden Markov-model (HMM) state of the windows. Blue colored points (low) indicate insignificant amount of introgression, while yellow-colored points (high) are outlier windows with significant introgression from the HMM analysis. Annotated genes in the outlier windows can be found in Table S3. TA = *Trichaptum abietinum* and TF = *T. fuscoviolaceum*. The figure is made in *R v4.0.2* using the packages *tidyverse* (Wickham et al. 2019) and *wesanderson* (Ram and Wickham, 2018).

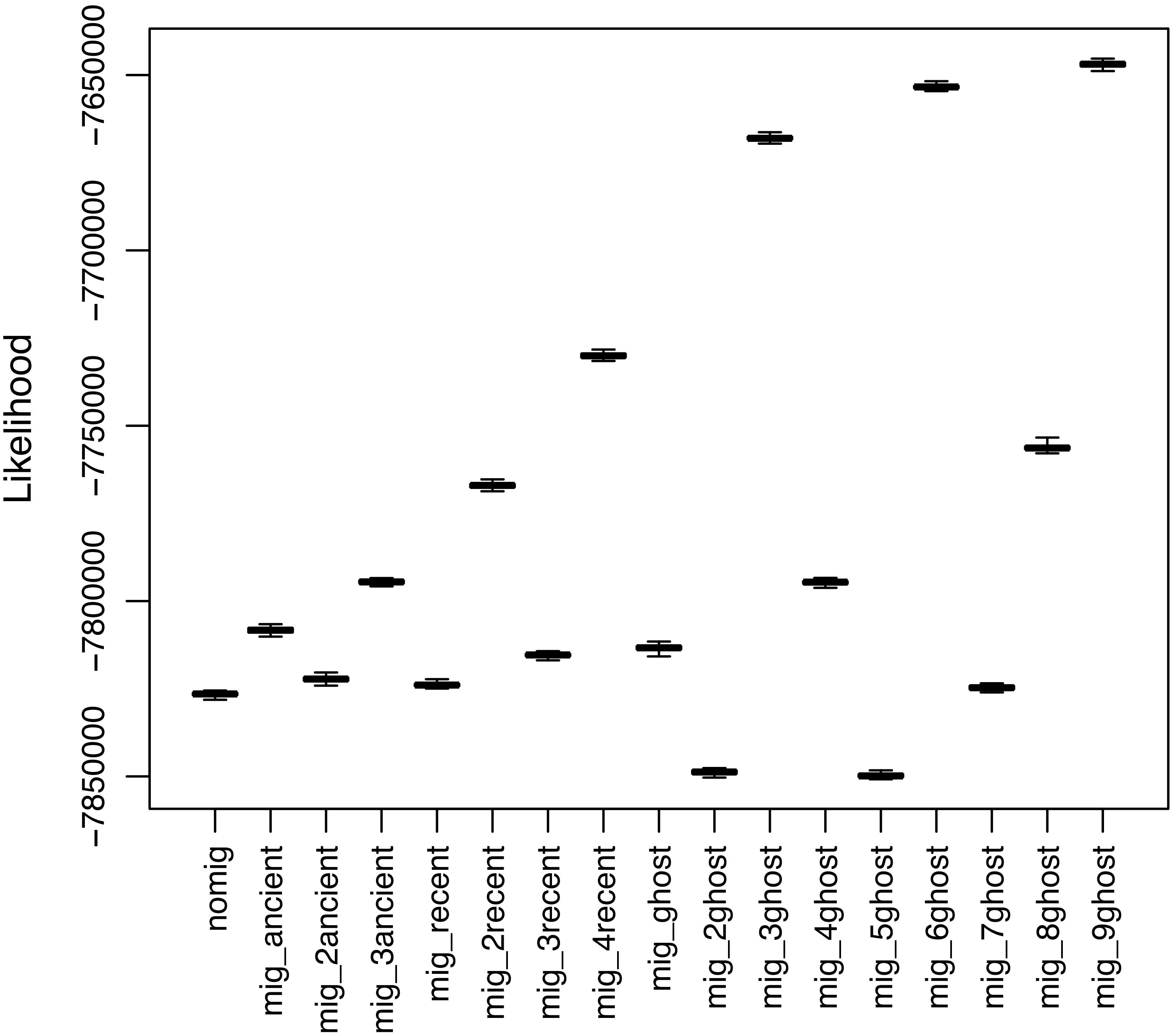


**Figure S11. The likelihood distributions support the best model based on AIC.** Likelihood distributions plotted for the different models tested in *fastsimcoal2* (Excoffier et al. 2021). The best model (mig_9ghost) does not have an overlapping distribution with the second best model (mig_6ghost). The distributions are plotted as boxplots with the likelihood on the y-axis and the name of the models on the x-axis. The model with the best likelihood (mig_9ghost) is also supported by AIC. See Figure S6 and Table S5 for further details on the models.

**Table S1. Overview of individuals collected and used for bioinformatic analyses.** The table includes information on name (collection ID), species designation (based on morphology, ITS sequence and Illumina sequence), collection site (area, longitude, latitude, and elevation), host substrate (substrate), name of collector and date of collection. TF = *Trichaptum fuscoviolaceum*, TA = *T. abietinum* and the outgroup TB = *T. biforme*.

| Collection ID | Species morphology | Species ITS | Species Illumina | Area | Latitude | Longitude | Elevation | Substrate | Collector | Collection date |
|---|---|---|---|---|---|---|---|---|---|---|
| TF-1000-1-M1 | *T. fuscoviolaceum* | *T. fuscoviolaceum* | *T. fuscoviolaceum* | CA, N.B., Charlotte County | 45.13741 N | 66.46785 W | 120 m | *Abies balsamea* | David Malloch | 09.10.2018 |
| TF-1002-2-M1 | *T. fuscoviolaceum* | *T. abietinum* | *T. abietinum* | CA, N.B., Charlotte County | 45.129722 N | 66.523889 W | 50 m | *Abies balsamea* | Inger Skrede & Dabao Lu | 09.10.2018 |
| TF-1002-3-M1 | *T. fuscoviolaceum* | *T. fuscoviolaceum* | *T. fuscoviolaceum* | CA, N.B., Charlotte County | 45.126944 N | 66.526944 W | 50 m | *Abies balsamea* | Inger Skrede & Dabao Lu | 09.10.2018 |
| TF-1002-4-M3 | *T. fuscoviolaceum* | *T. fuscoviolaceum* | *T. fuscoviolaceum* | CA, N.B., Charlotte County | 45.128611 N | 66.525000 W | 50 m | *Abies balsamea* | Inger Skrede & Dabao Lu | 09.10.2018 |
| TF-1002-5-M3 | *T. fuscoviolaceum* | *T. fuscoviolaceum* | *T. fuscoviolaceum* | CA, N.B., Charlotte County | 45.126944 N | 66.527222 W | 50 m | *Abies balsamea* | Inger Skrede & Dabao Lu | 09.10.2018 |
| TF-1002-7-M2 | *T. fuscoviolaceum* | *T. fuscoviolaceum* | *T. fuscoviolaceum* | CA, N.B., Charlotte County | 45.130833 N | 66.525000 W | 50 m | *Abies balsamea* | Inger Skrede & Dabao Lu | 09.10.2018 |
| TF-1002-10-M3 | *T. fuscoviolaceum* | *T. fuscoviolaceum* | *T. fuscoviolaceum* | CA, N.B., Charlotte County | 45.12927 N | 66.52469 W | 50 m | *Abies balsamea* | Amanda Bremner | 09.10.2018 |
| TF-1003-2-M1 | *T. fuscoviolaceum* | *T. fuscoviolaceum* | *T. fuscoviolaceum* | CA, N.B., Charlotte County | 45.169444 N | 66.459167 W | 10 m | *Picea rubens* | Inger Skrede & Dabao Lu | 09.10.2018 |
| TF-1003-3-M2 | *T. fuscoviolaceum* | *T. fuscoviolaceum* | *T. fuscoviolaceum* | CA, N.B., Charlotte County | 45.169167 N | 66.458889 W | 10 m | *Picea rubens* | Inger Skrede & Dabao Lu | 09.10.2018 |
| TF-1003-4-M2 | *T. fuscoviolaceum* | *T. fuscoviolaceum* | *T. fuscoviolaceum* | CA, N.B., Charlotte County | 45.173611 N | 66.465556 W | 10 m | *Abies balsamea* | Inger Skrede & Dabao Lu | 09.10.2018 |
| TF-1003-5-M2 | *T. fuscoviolaceum* | *T. fuscoviolaceum* | *T. fuscoviolaceum* | CA, N.B., Charlotte County | 45.168611 N | 66.461389 W | 10 m | *Picea rubens* | Inger Skrede & Dabao Lu | 09.10.2018 |
| TF-1004-1-M1 | *T. fuscoviolaceum* | *T. fuscoviolaceum* | *T. fuscoviolaceum* | CA, N.B., Sunbury County | 45.993333 N | 66.307500 W | 80 m | *Abies balsamea* | Inger Skrede & Dabao Lu | 10.10.2018 |
| TF-1004-3-M1 | *T. fuscoviolaceum* | *T. fuscoviolaceum* | *T. fuscoviolaceum* | CA, N.B., Sunbury County | 45.992222 N | 66.307222 W | 80 m | *Abies balsamea* | Inger Skrede & Dabao Lu | 10.10.2018 |

| | | | | | | | | | | |
|---|---|---|---|---|---|---|---|---|---|---|
| TF-1004-4-M2 | *T. fuscoviolaceum* | *T. fuscoviolaceum* | *T. fuscoviolaceum* | CA, N.B., Sunbury County | 45.992222 N | 66.307222 W | 80 m | *Abies balsamea* | Inger Skrede & Dabao Lu | 10.10.2018 |
| TF-1004-6-M2 | *T. fuscoviolaceum* | *T. fuscoviolaceum* | *T. fuscoviolaceum* | CA, N.B., Sunbury County | 46.00562 N | 66.40087 W | 130 m | *Abies balsamea* | Stephen R. Clayden | 10.10.2018 |
| TF-1004-6-dup-M2 | *T. fuscoviolaceum* | *T. fuscoviolaceum* | *T. fuscoviolaceum* | CA, N.B., Sunbury County | 45.991389 N | 66.307500 W | 80 m | *Abies balsamea* | Inger Skrede & Dabao Lu | 10.10.2018 |
| TF-1005-1-M1 | *T. fuscoviolaceum* | *T. abietinum* | *T. abietinum* | CA, N.B., Sunbury County | 46.035000 N | 66.325000 W | 130 m | *Picea mariana* | Inger Skrede & Dabao Lu | 10.10.2018 |
| TF-1007-1-M3 | *T. fuscoviolaceum* | *T. fuscoviolaceum* | *T. fuscoviolaceum* | CA, N.B., Gloucester County | 47.636944 N | 65.610833 W | 20 m | *Abies balsamea* | Inger Skrede & Dabao Lu | 11.10.2018 |
| TF-1007-2-M2 | *T. fuscoviolaceum* | *T. fuscoviolaceum* | *T. fuscoviolaceum* | CA, N.B., Gloucester County | 47.636944 N | 65.610833 W | 20 m | *Abies balsamea* | Inger Skrede & Dabao Lu | 11.10.2018 |
| TF-1009-1-M1 | *T. fuscoviolaceum* | *T. fuscoviolaceum* | *T. fuscoviolaceum* | CA, N.B., Restigouche County | 47.41611 N | 66.868056 W | 380 m | *Abies balsamea* | Inger Skrede & Dabao Lu | 12.10.2018 |
| TF-1009-2-M2 | *T. fuscoviolaceum* | *T. fuscoviolaceum* | *T. fuscoviolaceum* | CA, N.B., Restigouche County | 47.416111 N | 66.866944 W | 360 m | *Abies balsamea* | Inger Skrede & Dabao Lu | 12.10.2018 |
| TF-1009-3-M3 | *T. fuscoviolaceum* | *T. fuscoviolaceum* | *T. abietinum* | CA, N.B., Restigouche County | 47.418333 | 66.868889 W | 240 m | *Abies balsamea* | Inger Skrede & Dabao Lu | 12.10.2018 |
| TF-1009-4-M2 | *T. fuscoviolaceum* | *T. fuscoviolaceum* | *T. fuscoviolaceum* | CA, N.B., Restigouche County | 47.418056 N | 66.867778 W | 290 m | *Abies balsamea* | Inger Skrede & Dabao Lu | 12.10.2018 |
| TF-1011-1-M1 | *T. fuscoviolaceum* | *T. fuscoviolaceum* | *T. fuscoviolaceum* | CA, N.B., Victoria County | 46.894167 N | 67.398333 W | 170 m | *Abies balsamea* | Inger Skrede & Dabao Lu | 13.10.2018 |
| TF-1011-3-M1 | *T. fuscoviolaceum* | *T. fuscoviolaceum* | *T. fuscoviolaceum* | CA, N.B., Victoria County | 46.893333 N | 67.398333 W | 170 m | *Abies balsamea* | Inger Skrede & Dabao Lu | 13.10.2018 |
| TF-1011-4-M1 | *T. fuscoviolaceum* | *T. fuscoviolaceum* | *T. fuscoviolaceum* | CA, N.B., Victoria County | 46.892778 N | 67.398611 W | 170 m | *Abies balsamea* | Inger Skrede & Dabao Lu | 13.10.2018 |
| TF-1011-7-M2 | *T. fuscoviolaceum* | *T. fuscoviolaceum* | *T. fuscoviolaceum* | CA, N.B., Victoria County | 46.892500 N | 67.399167 W | 170 m | *Picea sp.* | Inger Skrede & Dabao Lu | 13.10.2018 |

| | | | | | | | | | | |
|---|---|---|---|---|---|---|---|---|---|---|
| TF-1011-8-M1 | *T. fuscoviolaceum* | *T. fuscoviolaceum* | *T. fuscoviolaceum* | CA, N.B., Victoria County | 46.902222 N | 67.400556 W | 170 m | *Abies balsamea* | Inger Skrede & Dabao Lu | 13.10.2018 |
| TF-1012-1-M2 | *T. fuscoviolaceum* | *T. fuscoviolaceum* | *T. fuscoviolaceum* | CA, N.B., Northumberland County | 46.783333 N | 66.516667 W | 390 m | *Abies balsamea* | Inger Skrede & Dabao Lu | 13.10.2018 |
| TF-1012-2-M1 | *T. fuscoviolaceum* | *T. fuscoviolaceum* | *T. fuscoviolaceum* | CA, N.B., Northumberland County | 46.783889 N | 66.525556 W | 390 m | *Abies balsamea* | Inger Skrede & Dabao Lu | 13.10.2018 |
| TF-1012-3-M3 | *T. fuscoviolaceum* | *T. fuscoviolaceum* | *T. fuscoviolaceum* | CA, N.B., Northumberland County | 46.784444 N | 66.526111 W | 400 m | *Abies balsamea* | Inger Skrede & Dabao Lu | 13.10.2018 |
| TF-1012-5-M2 | *T. fuscoviolaceum* | *T. fuscoviolaceum* | *T. fuscoviolaceum* | CA, N.B., Northumberland County | 46.785556 N | 60.525278 W | 380 m | *Abies balsamea* | Inger Skrede & Dabao Lu | 13.10.2018 |
| TF-1013-1-M2 | *T. fuscoviolaceum* | *T. fuscoviolaceum* | *T. fuscoviolaceum* | CA, N.B., York County | 45.9564 N | 66.6668 W | 50 m | *Abies balsamea* | Stephen R. Clayden | 12.10.2018 |
| TF-1013-2-M2 | *T. fuscoviolaceum* | *T. fuscoviolaceum* | *T. fuscoviolaceum* | CA, N.B., York County | 45.9564 N | 66.6668 W | 50 m | *Abies balsamea* | Stephen R. Clayden | 12.10.2018 |
| TF-1013-4-M2 | *T. fuscoviolaceum* | *T. fuscoviolaceum* | *T. abietinum* | CA, N.B., York County | 45.9564 N | 66.6668 W | 50 m | *Abies balsamea* | Stephen R. Clayden | 12.10.2018 |
| TF-1013-5-M2 | *T. fuscoviolaceum* | *T. fuscoviolaceum* | *T. fuscoviolaceum* | CA, N.B., York County | 45.9564 N | 66.6668 W | 50 m | *Abies balsamea* | Stephen R. Clayden | 12.10.2018 |
| TF-1013-8-M2 | *T. fuscoviolaceum* | *T. fuscoviolaceum* | *T. fuscoviolaceum* | CA, N.B., York County | 45.9564 N | 66.6668 W | 50 m | *Abies balsamea* | Stephen R. Clayden | 12.10.2018 |
| TF-1014-1-M2 | *T. fuscoviolaceum* | *T. fuscoviolaceum* | *T. fuscoviolaceum* | ITL, Pavia, Menconico | 44.80662 N | 9.31246 E | – | *Pinus nigra* | Carolina Girometta | 06.10.2018 |
| TF-1014-2-M1 | *T. fuscoviolaceum* | *T. fuscoviolaceum* | *T. fuscoviolaceum* | ITL, Pavia, Menconico | 44.80744 N | 9.31063 E | – | *Pinus nigra* | Carolina Girometta | 06.10.2018 |
| TF-1014-3-M1 | *T. fuscoviolaceum* | *T. fuscoviolaceum* | *T. fuscoviolaceum* | ITL, Pavia, Menconico | 44.80754 N | 9.31042 E | – | *Pinus nigra* | Carolina Girometta | 06.10.2018 |
| TF-1014-6-M3 | *T. fuscoviolaceum* | *T. fuscoviolaceum* | *T. fuscoviolaceum* | ITL, Pavia, Menconico | 44.81126 N | 9.30588 E | – | *Pinus nigra* | Carolina Girometta | 06.10.2018 |
| TF-1014-7-M1 | *T. fuscoviolaceum* | *T. fuscoviolaceum* | *T. fuscoviolaceum* | ITL, Pavia, Menconico | 44.81134 N | 9.30589 E | – | *Pinus nigra* | Carolina Girometta | 06.10.2018 |

| | | | | | | | | | | |
|---|---|---|---|---|---|---|---|---|---|---|
| TF-1014-7-M9 | *T. fuscoviolaceum* | *T. fuscoviolaceum* | *T. fuscoviolaceum* | ITL, Pavia, Menconico | 44.81134 N | 9.30589 E | – | *Pinus nigra* | Carolina Girometta | 06.10.2018 |
| TF-1014-8-M2 | *T. fuscoviolaceum* | *T. fuscoviolaceum* | *T. fuscoviolaceum* | ITL, Pavia, Menconico | 44.81137 N | 9.30661 E | – | *Pinus nigra* | Carolina Girometta | 06.10.2018 |
| TF-1014-9-M3 | *T. fuscoviolaceum* | *T. fuscoviolaceum* | *T. fuscoviolaceum* | ITL, Pavia, Menconico | 44.80795 N | 9.30954 E | – | *Pinus nigra* | Carolina Girometta | 06.10.2018 |
| TF-1014-10-M1 | *T. fuscoviolaceum* | *T. fuscoviolaceum* | *T. fuscoviolaceum* | ITL, Pavia, Menconico | 44.80520 N | 9.31298 E | – | *Pinus nigra* | Carolina Girometta | 06.10.2018 |
| TA-1002-8-M1 | *T. abietinum* | *T. abietinum* | *T. abietinum* | CA, N.B., Charlotte County | 45.129444 N | 66.524167 W | 50 m | *Abies balsamea* | Inger Skrede & Dabao Lu | 09.10.2018 |
| TA-1002-13-M2 | *T. abietinum* | *T. abietinum* | *T. abietinum* | CA, N.B., Charlotte County | 45.128611 N | 66.524444 W | 50 m | *Picea rubens* | Inger Skrede & Dabao Lu | 09.10.2018 |
| TA-1002-19-M1 | *T. abietinum* | *T. abietinum* | *T. abietinum* | CA, N.B., Charlotte County | 45.128611 N | 66.524722 W | 50 m | *Abies balsamea* | Inger Skrede & Dabao Lu | 09.10.2018 |
| TA-1002-27-M1 | *T. abietinum* | *T. abietinum* | *T. abietinum* | CA, N.B., Charlotte County | 45.128333 N | 66.526111 W | 50 m | *Picea rubens* | Inger Skrede & Dabao Lu | 09.10.2018 |
| TA-1002-35-M2 | *T. abietinum* | *T. abietinum* | *T. abietinum* | CA, N.B., Charlotte County | 45.128611 N | 66.526389 W | 50 m | *Picea rubens* | Inger Skrede & Dabao Lu | 09.10.2018 |
| TA-1003-1-M1 | *T. abietinum* | *T. abietinum* | *T. abietinum* | CA, N.B., Sunbury County | 45.991944 N | 66.306944 W | 60 m | *Abies balsamea* | Inger Skrede & Dabao Lu | 10.10.2018 |
| TA-1003-8-M1 | *T. abietinum* | *T. abietinum* | *T. abietinum* | CA, N.B., Sunbury County | 45.992778 N | 66.306944 W | 60 m | *Abies balsamea* | Inger Skrede & Dabao Lu | 10.10.2018 |
| TA-1003-17-M1 | *T. abietinum* | *T. abietinum* | *T. abietinum* | CA, N.B., Sunbury County | 45.990278 N | 66.306667 W | 60 m | *Abies balsamea* | Inger Skrede & Dabao Lu | 10.10.2018 |
| TA-1003-20-M2 | *T. abietinum* | *T. abietinum* | *T. abietinum* | CA, N.B., Sunbury County | 45.990278 N | 66.307222 W | 60 m | *Picea rubens* | Inger Skrede & Dabao Lu | 10.10.2018 |
| TA-1003-22-M2 | *T. abietinum* | *T. abietinum* | *T. abietinum* | CA, N.B., Sunbury County | 45.99056 N | 66.30690 W | 60 m | *Picea rubens* | Stepen R. Clayden | 10.10.2018 |
| TA-1007-1 | *T. abietinum* | *T. abietinum* | *T. abietinum* | CA, N.B., Gloucester County | 47.637500 N | 65.610000 W | 20 m | *Pinus cf. strobus* | Inger Skrede & Dabao Lu | 11.10.2018 |

| | | | | | | | | | | |
|---|---|---|---|---|---|---|---|---|---|---|
| TA-1007-3 | *T. abietinum* | *T. abietinum* | *T. abietinum* | CA, N.B., Gloucester County | 47.631389 N | 65.616111 W | 20 m | *Picea cf. glauca* | Inger Skrede & Dabao Lu | 11.10.2018 |
| TA-1007-5 | *T. abietinum* | *T. abietinum* | *T. abietinum* | CA, N.B., Gloucester County | 47.637500 N | 65.610556 W | 30 m | *Picea cf. glauca* | Inger Skrede & Dabao Lu | 11.10.2018 |
| TA-1007-6 | *T. abietinum* | *T. abietinum* | *T. abietinum* | CA, N.B., Gloucester County | 47.649722 N | 65.610833 W | 30 m | *Picea cf. glauca* | Inger Skrede & Dabao Lu | 11.10.2018 |
| TA-1007-17 | *T. abietinum* | *T. abietinum* | *T. abietinum* | CA, N.B., Gloucester County | 47.603333 N | 65.610833 W | 30 m | *Abies balsamea* | Inger Skrede & Dabao Lu | 11.10.2018 |
| TA-1009-1-M3 | *T. abietinum* | *T. abietinum* | *T. abietinum* | CA, N.B., Restigouche County | 47.418056 N | 66.866667 W | 240 m | *Picea cf. glauca* | Inger Skrede & Dabao Lu | 12.10.2018 |
| TA-1009-4-M1 | *T. abietinum* | *T. abietinum* | *T. abietinum* | CA, N.B., Restigouche County | 47.417778 N | 66.866944 W | 330 m | *Picea cf. glauca* | Inger Skrede & Dabao Lu | 12.10.2018 |
| TA-1009-8-M3 | *T. abietinum* | *T. abietinum* | *T. abietinum* | CA, N.B., Restigouche County | 47.416944 N | 66.867222 W | 330 m | *Picea cf. glauca* | Inger Skrede & Dabao Lu | 12.10.2018 |
| TA-1009-12-M1 | *T. abietinum* | *T. abietinum* | *T. abietinum* | CA, N.B., Restigouche County | 47.417778 N | 66.866111 W | 340 m | *Picea sp.* | Inger Skrede & Dabao Lu | 12.10.2018 |
| TA-1009-19-M3 | *T. abietinum* | *T. abietinum* | *T. abietinum* | CA, N.B., Restigouche County | 47.418611 N | 66.881389 W | 220 m | *Picea sp* | Inger Skrede & Dabao Lu | 12.10.2018 |
| TA-1011-12-M2 | *T. abietinum* | *T. abietinum* | *T. abietinum* | CA, N.B., Victoria County | 46.894167 N | 67.398333 W | 170 m | *Picea cf. rubens* | Inger Skrede & Dabao Lu | 13.10.2018 |
| TA-1011-19-M1 | *T. abietinum* | *T. abietinum* | *T. abietinum* | CA, N.B., Victoria County | 46.892500 N | 67.399444 W | 170 m | *Picea cf. rubens* | Inger Skrede & Dabao Lu | 13.10.2018 |
| TA-1011-23-M1 | *T. abietinum* | *T. abietinum* | *T. abietinum* | CA, N.B., Victoria County | 46.900833 N | 67.400278 W | 170 m | *Picea sp.* | Inger Skrede & Dabao Lu | 13.10.2018 |
| TA-1011-26-M1 | *T. abietinum* | *T. abietinum* | *T. abietinum* | CA, N.B., Victoria County | 46.893611 N | 67.400000 W | 170 m | *Abies balsamea* | Inger Skrede & Dabao Lu | 13.10.2018 |
| TA-1011-31-M1 | *T. abietinum* | *T. abietinum* | *T. abietinum* | CA, N.B., Victoria County | 46.893333 N | 67.401111 W | 170 m | *Abies balsamea* | Inger Skrede & Dabao Lu | 13.10.2018 |

| | | | | | | | | | | |
|---|---|---|---|---|---|---|---|---|---|---|
| TA-1012-3-M1 | *T. abietinum* | *T. abietinum* | *T. abietinum* | CA, N.B., Northumberland County | 46.783889 N | 66.525556 W | 390 m | *Picea rubens* | Inger Skrede & Dabao Lu | 13.10.2018 |
| TA-1012-5-M1 | *T. abietinum* | *T. abietinum* | *T. abietinum* | CA, N.B., Northumberland County | 46.784167 N | 66.059444 W | 390 m | *Abies balsamea* | Inger Skrede & Dabao Lu | 13.10.2018 |
| TA-1012-7-M1 | *T. abietinum* | *T. abietinum* | *T. abietinum* | CA, N.B., Northumberland County | 46.784167 N | 66.059444 W | 390 m | *Picea rubens* | Inger Skrede & Dabao Lu | 13.10.2018 |
| TA-1012-11-M1 | *T. abietinum* | *T. abietinum* | *T. abietinum* | CA, N.B., Northumberland County | 46.785556 N | 66.525556 W | 390 m | *Picea rubens* | Inger Skrede & Dabao Lu | 13.10.2018 |
| TA-1012-17-M1 | *T. abietinum* | *T. abietinum* | *T. abietinum* | CA, N.B., Northumberland County | 46.784722 N | 66.524722 W | 380 m | *Picea sp.* | Inger Skrede & Dabao Lu | 13.10.2018 |
| TA-1013-3-M2 | *T. abietinum* | *T. abietinum* | *T. abietinum* | CA, N.B., York County | 45.9564 N | 66.6668 W | 50 m | *Abies balsamea* | Stephen R. Clayden | 12.10.2018 |
| TA-1013-4-M1 | *T. abietinum* | *T. abietinum* | *T. abietinum* | CA, N.B., York County | 45.9564 N | 66.6668 W | 50 m | *Tsuga canadensis* | Stephen R. Clayden | 12.10.2018 |
| TA-1013-5-M1 | *T. abietinum* | *T. abietinum* | *T. abietinum* | CA, N.B., York County | 45.9564 N | 66.6668 W | 50 m | *Tsuga canadensis* | Stephen R. Clayden | 12.10.2018 |
| TA-1013-7-M1 | *T. abietinum* | *T. abietinum* | *T. abietinum* | CA, N.B., York County | 45.9564 N | 66.6668 W | 50 m | *Picea rubens* | Stephen R. Clayden | 12.10.2018 |
| TA-1013-9-M1 | *T. abietinum* | *T. abietinum* | *T. abietinum* | CA, N.B., York County | 45.9564 N | 66.6668 W | 50 m | *Picea rubens* | Stephen R. Clayden | 12.10.2018 |
| TB-1013-1-M2 | *T. fuscoviolaceum* | *T. biforme* | *T. biforme* | CA, N. B., York County | 45.9564 N | 66.6668 W | 50 m | *Abies balsamea* | Stephen R. Clayden | 12.10.2018 |

**Table S2, *Trichaptum fuscoviolaceum* individuals mated as predicted, while *T. abietinum* crossed with T. fuscoviolaceum individuals did not.** The table includes cross name (Cross ID), monokaryotic individuals crossed (Mate pairs), mating loci differences and similarities between the crosses (Mating type (MAT)), populations crossed (Populations), expected outcome (Prediction) and actual outcome by observation (Yes) or no observation (No) of clamp connections (Clamp).

| ***T. abietinum* × *T. fuscoviolaceum*** | | | | | |
|---|---|---|---|---|---|
| **Cross ID** | **Mate pairs** | **Mating type (*MAT*)** | **Populations** | **Prediction** | **Clamp** |
| TFTAX1 | TF10147M9 × TA10264M3 | Ident. *MATA*, dist. *MATB* | It × Eu | Incompatible | No |
| TFTAX2 | TF10147M9 × TA10058M1 | Ident. *MATA*, dist. *MATB* | It × NAmA | Incompatible | No |
| TFTAX3 | TF101410M1 × TA10355M3 | Dist. *MATA*, ident. *MATB* | It × Eu | Incompatible | No |
| TFTAX4 | TF10141M2 × TA10139M1 | Dist. *MATA*, ident. *MATB* | It × NAmB | Incompatible | No |
| TFTAX5 | TF101410M1 × TA10264M3 | Dist. *MAT*s | It × Eu | Compatible | No |
| TFTAX6 | TF101410M1 × TA10139M1 | Dist. *MAT*s | It × NAmB | Compatible | No |
| TFTAX7 | TF101410M1 × TA10058M1 | Dist. *MAT*s | It × NAmA | Compatible | No |
| TFTAX8 | TF10034M2 × TA10139M1 | Dist. *MAT*s | Can × NAmB | Compatible | No |
| TFTAX9 | TF10034M2 × TA10264M3 | Dist. *MAT*s | Can × Eu | Compatible | No |
| TFTAX11 | TF10034M2 × TA10058M1 | Dist. *MAT*s | Can × NAmA | Compatible | No |
| ***T. fuscoviolaceum* × *T. fuscoviolaceum*** | | | | | |
| **Cross ID** | **Mate pairs** | **Mating type (*MAT*)** | **Populations** | **Prediction** | **Clamp** |
| TFX1 | TF10147M1 × TF10147M1 | Ident. *MAT* | It × It | Incompatible | No |
| TFX2 | TF10147M1 × TF10147M9 | Dist. α *MATA* and *MATB* | It × It | Compatible | Yes |
| TFX3 | TF10147M1 × TF10143M3 | Dist. α *MATA,* ident. *MATB* | It × It | Incompatible | No |
| TFX4 | TF10147M9 × TF10141M2 | Dist. β *MATA* and *MATB* | It × It | Compatible | Yes |
| TFX5 | TF10032M1 × TF10135M2 | Dist. β *MATA* and *MATB* | Can × Can | Compatible | Yes |
| TFX7 | TF10091M1 × TF10135M2 | Dist. α *MAT* and *MATB* | Can × Can | Compatible | Yes |
| TFX9 | TF10034M2 × TF10091M1 | Dist. *MATA*, ident. *MATB* | Can × Can | Incompatible | No |
| TFX10 | TF10122M1 × TF10147M1 | Dist. β *MATA* and *MATB* | It × Can | Compatible | Yes |
| TFX11 | TF101410M1 × TF10122M1 | Dist. α *MATA* and *MATB* | It × Can | Compatible | Yes |
| TFX12 | TF10141M2 × TF10122M1 | Dist. *MATA*, ident. *MATB* | It × Can | Incompatible | No |

Dist. = distinct, ident. = identical, TA = *Trichaptum abietinum*, TF = *T. fuscoviolaceum*, It = Italian, Can = Canadian, Eu = European, NAmA = North American A, NAmB = North American B

**Table S3. An overview of the Hidden Markov-model outliers from the $f_{dM}$ genome scan.** The headers indicate the phylogenetic typology of the test; (((P1, P2), P3), O), where P1, P2 and P3 are populations investigated for introgression and O is the outgroup. A positive $f_{dM}$ value indicates more shared derived polymorphisms than expected between P2 and P3, while a negative value indicates more shared derived polymorphisms than expected between P1 and P3. The table also denotes which scaffold the genes are in, including the start and end of the gene on that scaffold, how many sites that were used to estimate the $f_{dM}$ value in the specific window (Sites used), the name of the genes from the annotated genome (Gene), and a note on the function of the genes. TA = *Trichaptum abietinum* and TF = *T. fuscoviolaceum*.

| (((North American B TA, North American A TA), Canadian TF), *T. biforme*) | | | | | | |
|---|---|---|---|---|---|---|
| **Scaffold** | **Start** | **End** | **Sites used** | $f_{dM}$ | **Gene** | **Note** |
| Scaffold01 | 745385 | 745484 | 122 | 0.1053 | trnascan-Scaffold01-noncoding-Thr_TGT-gene-7.19 | Protein of unknown function |
| Scaffold01 | 745617 | 746235 | 122 | 0.1053 | maker-Scaffold01-snap-gene-7.9 | Protein of unknown function |
| Scaffold01 | 746860 | 752545 | 122 | 0.1053 | snap_masked-Scaffold01-processed-gene-7.3 | Similar to ATP1A1: Sodium/potassium-transporting ATPase subunit alpha-1 (Equus caballus OX=9796) |
| Scaffold01 | 752671 | 755023 | 122 | 0.1053 | maker-Scaffold01-snap-gene-7.12 | Protein of unknown function |
| Scaffold01 | 900331 | 900881 | 133 | 0.1251 | maker-Scaffold01-snap-gene-9.2 | Protein of unknown function |
| Scaffold01 | 901468 | 902851 | 133 | 0.1251 | snap_masked-Scaffold01-processed-gene-9.31 | Protein of unknown function |
| Scaffold01 | 902938 | 905385 | 133 | 0.1251 | snap_masked-Scaffold01-processed-gene-9.39 | Protein of unknown function |
| Scaffold01 | 906260 | 907088 | 133 | 0.1251 | maker-Scaffold01-exonerate_protein2genome-gene-9.29 | Protein of unknown function |
| Scaffold01 | 909092 | 910780 | 133 | 0.1251 | maker-Scaffold01-snap-gene-9.4 | Protein of unknown function |
| Scaffold01 | 912091 | 912422 | 133 | 0.1251 | snap_masked-Scaffold01-processed-gene-9.40 | Similar to EMC4: ER membrane protein complex subunit 4 (Saccharomyces cerevisiae (strain ATCC 204508 / S288c) OX=559292) |
| Scaffold01 | 913710 | 915599 | 133 | 0.1251 | maker-Scaffold01-snap-gene-9.5 | Similar to COX17: Cytochrome c oxidase copper chaperone (Homo sapiens OX=9606) |
| Scaffold01 | 915247 | 917294 | 133 | 0.1251 | maker-Scaffold01-snap-gene-9.16 | Similar to GST: Glutathione S-transferase (Plasmodium vivax OX=5855) |
| Scaffold01 | 918913 | 919817 | 133 | 0.1251 | snap_masked-Scaffold01-abinit-gene-9.25 | Protein of unknown function |
| Scaffold01 | 2975369 | 2975485 | 137 | 0.1062 | maker-Scaffold01-exonerate_protein2genome-gene-29.152 | Protein of unknown function |

| | | | | | | |
|---|---|---|---|---|---|---|
| Scaffold03 | 1384980 | 1386656 | 110 | 0.1063 | genemark-Scaffold03-processed-gene-13.17 | Similar to NEP1: Ribosomal RNA small subunit methyltransferase NEP1 (Candida albicans OX=5476) |
| Scaffold03 | 1387398 | 1388969 | 110 | 0.1063 | maker-Scaffold03-snap-gene-13.15 | Similar to fmdA: Formamidase (Methylophilus methylotrophus OX=17) |
| Scaffold03 | 1391654 | 1393548 | 110 | 0.1063 | maker-Scaffold03-snap-gene-14.40 | Protein of unknown function |
| Scaffold03 | 1620126 | 1622452 | 106 | 0.1538 | genemark-Scaffold03-processed-gene-16.4 | Similar to GRC3: Polynucleotide 5'-hydroxyl-kinase GRC3 (Cryptococcus neoformans var. neoformans serotype D (strain JEC21 / ATCC MYA-565) OX=214684) |
| Scaffold03 | 1624233 | 1626184 | 106 | 0.1538 | maker-Scaffold03-snap-gene-16.42 | Similar to DAL1: Allantoinase (Saccharomyces cerevisiae (strain ATCC 204508 / S288c) OX=559292) |
| Scaffold03 | 1626282 | 1628466 | 106 | 0.1538 | maker-Scaffold03-snap-gene-16.43 | Protein of unknown function |
| Scaffold03 | 1628643 | 1630494 | 106 | 0.1538 | maker-Scaffold03-snap-gene-16.53 | Protein of unknown function |
| Scaffold03 | 1631588 | 1632628 | 106 | 0.1538 | maker-Scaffold03-snap-gene-16.54 | Protein of unknown function |
| Scaffold03 | 1632377 | 1632573 | 106 | 0.1538 | maker-Scaffold03-exonerate_est2genome-gene-16.2 | Protein of unknown function |
| Scaffold03 | 1632645 | 1632758 | 106 | 0.1538 | maker-Scaffold03-exonerate_protein2genome-gene-16.47 | Protein of unknown function |
| Scaffold03 | 1633221 | 1636431 | 106 | 0.1538 | maker-Scaffold03-exonerate_protein2genome-gene-16.48 | Similar to RDR1: Probable RNA-dependent RNA polymerase 1 (Oryza sativa subsp. japonica OX=39947) |
| Scaffold03 | 1637269 | 1638202 | 106 | 0.1538 | maker-Scaffold03-exonerate_protein2genome-gene-16.56 | Similar to SEC14: SEC14 cytosolic factor (Saccharomyces cerevisiae (strain ATCC 204508 / S288c) OX=559292) |
| Scaffold05 | 2389277 | 2393966 | 111 | 0.1137 | maker-Scaffold05-snap-gene-24.8 | Protein of unknown function |
| Scaffold05 | 2480001 | 2500000 | 125 | 0.1398 | No annotated genes | No annotated genes |
| Scaffold06 | 1741847 | 1744120 | 353 | 0.104 | maker-Scaffold06-snap-gene-17.52 | Similar to obg: GTPase Obg (Rippkaea orientalis (strain PCC 8801) OX=41431) |
| Scaffold06 | 1745220 | 1746591 | 353 | 0.104 | snap_masked-Scaffold06-processed-gene-17.7 | Similar to cel1: Cellulose-growth-specific protein (Agaricus bisporus OX=5341) |

| | | | | | | |
|---|---|---|---|---|---|---|
| Scaffold06 | 1747150 | 1751861 | 353 | 0.104 | maker-Scaffold06-snap-gene-17.53 | Similar to fgenesh1_kg.2_#_1379_#_Locus12621v1rpkm4.09: 4-O-methyltransferase 1 (Phanerochaete chrysosporium (strain RP-78 / ATCC MYA-4764 / FGSC 9002) OX=273507) |
| Scaffold06 | 1752774 | 1753281 | 353 | 0.104 | maker-Scaffold06-exonerate_protein2genome-gene-17.176 | Protein of unknown function |
| Scaffold06 | 1753385 | 1755997 | 353 | 0.104 | maker-Scaffold06-exonerate_protein2genome-gene-17.72 | Protein of unknown function |
| Scaffold06 | 1756135 | 1759294 | 353 | 0.104 | snap_masked-Scaffold06-processed-gene-17.23 | Similar to SPBC530.05: Uncharacterized transcriptional regulatory protein C530.05 (Schizosaccharomyces pombe (strain 972 / ATCC 24843) OX=284812) |
| Scaffold06 | 1759332 | 1759612 | 353 | 0.104 | snap_masked-Scaffold06-processed-gene-17.24 | Protein of unknown function |
| Scaffold07 | 3840869 | 3842531 | 163 | 0.1162 | maker-Scaffold07-snap-gene-38.79 | Similar to ZFAND2A: AN1-type zinc finger protein 2A (Pongo abelii OX=9601) |
| Scaffold07 | 3843021 | 3844141 | 163 | 0.1162 | snap_masked-Scaffold07-abinit-gene-38.26 | Protein of unknown function |
| Scaffold07 | 3845137 | 3848851 | 163 | 0.1162 | maker-Scaffold07-snap-gene-38.49 | Similar to PHB2: Prohibitin-2 (Saccharomyces cerevisiae (strain ATCC 204508 / S288c) OX=559292) |
| Scaffold07 | 3848870 | 3850190 | 163 | 0.1162 | maker-Scaffold07-exonerate_protein2genome-gene-38.89 | Protein of unknown function |
| Scaffold07 | 3850825 | 3853362 | 163 | 0.1162 | maker-Scaffold07-snap-gene-38.40 | Similar to exosc3: Putative exosome complex component rrp40 (Dictyostelium discoideum OX=44689) |
| Scaffold07 | 3853955 | 3856502 | 163 | 0.1162 | maker-Scaffold07-snap-gene-38.51 | Protein of unknown function |
| Scaffold07 | 3857642 | 3858990 | 163 | 0.1162 | maker-Scaffold07-snap-gene-38.52 | Protein of unknown function |
| Scaffold08 | 987415 | 988459 | 107 | 0.1363 | maker-Scaffold08-snap-gene-9.25 | Protein of unknown function |
| Scaffold08 | 990031 | 990293 | 107 | 0.1363 | maker-Scaffold08-exonerate_protein2genome-gene-10.96 | Protein of unknown function |
| Scaffold08 | 990859 | 996481 | 107 | 0.1363 | maker-Scaffold08-snap-gene-10.13 | Protein of unknown function |
| Scaffold08 | 996522 | 997428 | 107 | 0.1363 | maker-Scaffold08-snap-gene-10.18 | Protein of unknown function |
| Scaffold08 | 997701 | 997814 | 107 | 0.1363 | trnascan-Scaffold08-noncoding-Gly_GCC-gene-10.66 | Protein of unknown function |

| | | | | | | |
|---|---|---|---|---|---|---|
| Scaffold08 | 998636 | 999746 | 107 | 0.1363 | maker-Scaffold08-exonerate_protein2genome-gene-10.16 | Similar to ECI3: Enoyl-CoA delta isomerase 3 (Arabidopsis thaliana OX=3702) |
| Scaffold09 | 2701803 | 2709281 | 146 | 0.1067 | maker-Scaffold09-augustus-gene-27.6 | Similar to KES1: Protein KES1 (Ustilago maydis (strain 521 / FGSC 9021) OX=237631) |
| Scaffold09 | 2709464 | 2710425 | 146 | 0.1067 | maker-Scaffold09-augustus-gene-27.1 | Protein of unknown function |
| Scaffold09 | 2711618 | 2713971 | 146 | 0.1067 | maker-Scaffold09-snap-gene-27.54 | Similar to can: Carbonic anhydrase 2 (Shigella flexneri OX=623) |
| Scaffold10 | 1740426 | 1741739 | 222 | 0.1555 | snap_masked-Scaffold10-abinit-gene-17.14 | Protein of unknown function |
| Scaffold10 | 1742972 | 1744147 | 222 | 0.1555 | snap_masked-Scaffold10-processed-gene-17.14 | Protein of unknown function |
| Scaffold10 | 1744941 | 1746814 | 222 | 0.1555 | snap_masked-Scaffold10-processed-gene-17.15 | Similar to MEL: Alpha-galactosidase (Saccharomyces mikatae OX=114525) |
| Scaffold11 | 2845048 | 2846438 | 110 | 0.1194 | maker-Scaffold11-snap-gene-28.58 | Similar to SLC25A17: Peroxisomal membrane protein PMP34 (Homo sapiens OX=9606) |
| Scaffold11 | 2846866 | 2850623 | 110 | 0.1194 | genemark-Scaffold11-processed-gene-28.8 | Protein of unknown function |
| **(((North American B TA, North American A TA), Italian TF), *T. biforme*)** | | | | | | |
| **Scaffold** | **Start** | **End** | **Sites used** | $f_{dM}$ | **Gene** | **Note** |
| Scaffold01 | 900331 | 900881 | 153 | 0.1181 | maker-Scaffold01-snap-gene-9.2 | Protein of unknown function |
| Scaffold01 | 901468 | 902851 | 153 | 0.1181 | snap_masked-Scaffold01-processed-gene-9.31 | Protein of unknown function |
| Scaffold01 | 902938 | 905385 | 153 | 0.1181 | snap_masked-Scaffold01-processed-gene-9.39 | Protein of unknown function |
| Scaffold01 | 906260 | 907088 | 153 | 0.1181 | maker-Scaffold01-exonerate_protein2genome-gene-9.29 | Protein of unknown function |
| Scaffold01 | 909092 | 910780 | 153 | 0.1181 | maker-Scaffold01-snap-gene-9.4 | Protein of unknown function |
| Scaffold01 | 912091 | 912422 | 153 | 0.1181 | snap_masked-Scaffold01-processed-gene-9.40 | Similar to EMC4: ER membrane protein complex subunit 4 (Saccharomyces cerevisiae (strain ATCC 204508 / S288c) OX=559292) |
| Scaffold01 | 913710 | 915599 | 153 | 0.1181 | maker-Scaffold01-snap-gene-9.5 | Similar to COX17: Cytochrome c oxidase copper chaperone (Homo sapiens OX=9606) |

| | | | | | | |
|---|---|---|---|---|---|---|
| Scaffold01 | 915247 | 917294 | 153 | 0.1181 | maker-Scaffold01-snap-gene-9.16 | Similar to GST: Glutathione S-transferase (Plasmodium vivax OX=5855) |
| Scaffold01 | 918913 | 919817 | 153 | 0.1181 | snap_masked-Scaffold01-abinit-gene-9.25 | Protein of unknown function |
| Scaffold01 | 2975369 | 2975485 | 145 | 0.1387 | maker-Scaffold01-exonerate_protein2genome-gene-29.152 | Protein of unknown function |
| Scaffold02 | 4983917 | 4984611 | 104 | 0.1167 | maker-Scaffold02-snap-gene-50.56 | Protein of unknown function |
| Scaffold02 | 4986952 | 4987539 | 104 | 0.1167 | maker-Scaffold02-exonerate_protein2genome-gene-49.140 | Protein of unknown function |
| Scaffold02 | 4989157 | 4990010 | 104 | 0.1167 | maker-Scaffold02-exonerate_protein2genome-gene-50.0 | Similar to imp1: Mitochondrial inner membrane protease subunit 1 (Schizosaccharomyces pombe (strain 972 / ATCC 24843) OX=284812) |
| Scaffold02 | 4991580 | 4993084 | 104 | 0.1167 | snap_masked-Scaffold02-processed-gene-50.18 | Similar to TTC1: Tetratricopeptide repeat protein 1 (Bos taurus OX=9913) |
| Scaffold02 | 4993236 | 4994391 | 104 | 0.1167 | snap_masked-Scaffold02-processed-gene-50.3 | Similar to dltE: Uncharacterized oxidoreductase DltE (Bacillus subtilis (strain 168) OX=224308) |
| Scaffold02 | 4994792 | 4998647 | 104 | 0.1167 | maker-Scaffold02-snap-gene-50.59 | Similar to bglX: Periplasmic beta-glucosidase (Escherichia coli (strain K12) OX=83333) |
| Scaffold03 | 1620126 | 1622452 | 117 | 0.1619 | genemark-Scaffold03-processed-gene-16.4 | Similar to GRC3: Polynucleotide 5'-hydroxyl-kinase GRC3 (Cryptococcus neoformans var. neoformans serotype D (strain JEC21 / ATCC MYA-565) OX=214684) |
| Scaffold03 | 1624233 | 1626184 | 117 | 0.1619 | maker-Scaffold03-snap-gene-16.42 | Similar to DAL1: Allantoinase (Saccharomyces cerevisiae (strain ATCC 204508 / S288c) OX=559292) |
| Scaffold03 | 1626282 | 1628466 | 117 | 0.1619 | maker-Scaffold03-snap-gene-16.43 | Protein of unknown function |
| Scaffold03 | 1628643 | 1630494 | 117 | 0.1619 | maker-Scaffold03-snap-gene-16.53 | Protein of unknown function |
| Scaffold03 | 1631588 | 1632628 | 117 | 0.1619 | maker-Scaffold03-snap-gene-16.54 | Protein of unknown function |
| Scaffold03 | 1632377 | 1632573 | 117 | 0.1619 | maker-Scaffold03-exonerate_est2genome-gene-16.2 | Protein of unknown function |
| Scaffold03 | 1632645 | 1632758 | 117 | 0.1619 | maker-Scaffold03-exonerate_protein2genome-gene-16.47 | Protein of unknown function |

| | | | | | | |
|---|---|---|---|---|---|---|
| Scaffold03 | 1633221 | 1636431 | 117 | 0.1619 | maker-Scaffold03-exonerate_protein2genome-gene-16.48 | Similar to RDR1: Probable RNA-dependent RNA polymerase 1 (Oryza sativa subsp. japonica OX=39947) |
| Scaffold03 | 1637269 | 1638202 | 117 | 0.1619 | maker-Scaffold03-exonerate_protein2genome-gene-16.56 | Similar to SEC14: SEC14 cytosolic factor (Saccharomyces cerevisiae (strain ATCC 204508 / S288c) OX=559292) |
| Scaffold05 | 701713 | 703798 | 153 | 0.1408 | maker-Scaffold05-exonerate_protein2genome-gene-7.20 | Similar to pik-1: Pelle-like serine/threonine-protein kinase pik-1 (Caenorhabditis elegans OX=6239) |
| Scaffold05 | 704321 | 707167 | 153 | 0.1408 | maker-Scaffold05-snap-gene-7.2 | Similar to STY8: Serine/threonine-protein kinase STY8 (Arabidopsis thaliana OX=3702) |
| Scaffold05 | 707896 | 709998 | 153 | 0.1408 | maker-Scaffold05-snap-gene-7.3 | Similar to splB: Dual specificity protein kinase splB (Dictyostelium discoideum OX=44689) |
| Scaffold05 | 710120 | 711718 | 153 | 0.1408 | maker-Scaffold05-snap-gene-7.16 | Similar to SPAC4A8.06c: AB hydrolase superfamily protein C4A8.06c (Schizosaccharomyces pombe (strain 972 / ATCC 24843) OX=284812) |
| Scaffold05 | 712158 | 713655 | 153 | 0.1408 | maker-Scaffold05-exonerate_protein2genome-gene-7.31 | Similar to SPAC5D6.12: Uncharacterized protein C5D6.12 (Schizosaccharomyces pombe (strain 972 / ATCC 24843) OX=284812) |
| Scaffold05 | 713908 | 718057 | 153 | 0.1408 | maker-Scaffold05-snap-gene-7.17 | Similar to ARHGAP39: Rho GTPase-activating protein 39 (Homo sapiens OX=9606) |
| Scaffold05 | 718805 | 719236 | 153 | 0.1408 | maker-Scaffold05-exonerate_protein2genome-gene-7.39 | Protein of unknown function |
| Scaffold05 | 2480001 | 2500000 | 122 | 0.1628 | No annotated genes | No annotated genes |
| Scaffold07 | 3840869 | 3842531 | 167 | 0.1483 | maker-Scaffold07-snap-gene-38.79 | Similar to ZFAND2A: AN1-type zinc finger protein 2A (Pongo abelii OX=9601) |
| Scaffold07 | 3843021 | 3844141 | 167 | 0.1483 | snap_masked-Scaffold07-abinit-gene-38.26 | Protein of unknown function |
| Scaffold07 | 3845137 | 3848851 | 167 | 0.1483 | maker-Scaffold07-snap-gene-38.49 | Similar to PHB2: Prohibitin-2 (Saccharomyces cerevisiae (strain ATCC 204508 / S288c) OX=559292) |
| Scaffold07 | 3848870 | 3850190 | 167 | 0.1483 | maker-Scaffold07-exonerate_protein2genome-gene-38.89 | Protein of unknown function |

| | | | | | | |
|---|---|---|---|---|---|---|
| Scaffold07 | 3850825 | 3853362 | 167 | 0.1483 | maker-Scaffold07-snap-gene-38.40 | Similar to exosc3: Putative exosome complex component rrp40 (Dictyostelium discoideum OX=44689) |
| Scaffold07 | 3853955 | 3856502 | 167 | 0.1483 | maker-Scaffold07-snap-gene-38.51 | Protein of unknown function |
| Scaffold07 | 3857642 | 3858990 | 167 | 0.1483 | maker-Scaffold07-snap-gene-38.52 | Protein of unknown function |
| Scaffold08 | 987415 | 988459 | 111 | 0.1333 | maker-Scaffold08-snap-gene-9.25 | Protein of unknown function |
| Scaffold08 | 990031 | 990293 | 111 | 0.1333 | maker-Scaffold08-exonerate_protein2genome-gene-10.96 | Protein of unknown function |
| Scaffold08 | 990859 | 996481 | 111 | 0.1333 | maker-Scaffold08-snap-gene-10.13 | Protein of unknown function |
| Scaffold08 | 996522 | 997428 | 111 | 0.1333 | maker-Scaffold08-snap-gene-10.18 | Protein of unknown function |
| Scaffold08 | 997701 | 997814 | 111 | 0.1333 | trnascan-Scaffold08-noncoding-Gly_GCC-gene-10.66 | Protein of unknown function |
| Scaffold08 | 998636 | 999746 | 111 | 0.1333 | maker-Scaffold08-exonerate_protein2genome-gene-10.16 | Similar to ECI3: Enoyl-CoA delta isomerase 3 (Arabidopsis thaliana OX=3702) |
| Scaffold09 | 3641160 | 3641384 | 185 | 0.1245 | maker-Scaffold09-exonerate_protein2genome-gene-36.137 | Protein of unknown function |
| Scaffold09 | 3642465 | 3643793 | 185 | 0.1245 | maker-Scaffold09-snap-gene-36.38 | Similar to truC: tRNA pseudouridine synthase C (Yersinia pestis OX=632) |
| Scaffold09 | 3643841 | 3652490 | 185 | 0.1245 | maker-Scaffold09-snap-gene-36.30 | Similar to PPO1: Polyphenol oxidase 1 (Agaricus bisporus OX=5341) |
| Scaffold09 | 3652622 | 3655673 | 185 | 0.1245 | maker-Scaffold09-exonerate_protein2genome-gene-36.142 | Similar to pr1: Aspartic protease (Phaffia rhodozyma OX=5421) |
| Scaffold09 | 3657013 | 3657605 | 185 | 0.1245 | maker-Scaffold09-exonerate_protein2genome-gene-36.152 | Protein of unknown function |
| Scaffold09 | 3658494 | 3659856 | 185 | 0.1245 | maker-Scaffold09-snap-gene-36.41 | Similar to SPBC216.03: UPF0659 protein C216.03 (Schizosaccharomyces pombe (strain 972 / ATCC 24843) OX=284812) |
| Scaffold10 | 1740426 | 1741739 | 241 | 0.1656 | snap_masked-Scaffold10-abinit-gene-17.14 | Protein of unknown function |
| Scaffold10 | 1742972 | 1744147 | 241 | 0.1656 | snap_masked-Scaffold10-processed-gene-17.14 | Protein of unknown function |
| Scaffold11 | 2845048 | 2846438 | 113 | 0.1259 | maker-Scaffold11-snap-gene-28.58 | Similar to SLC25A17: Peroxisomal membrane protein PMP34 (Homo sapiens OX=9606) |
| Scaffold11 | 2846866 | 2850623 | 113 | 0.1259 | genemark-Scaffold11-processed-gene-28.8 | Protein of unknown function |

**Table S4. Overview of which cellular processes the outlier genes from the $f_{dM}$ genome scan are involved in, according to the GO enrichment analysis.** The headers indicate the phylogenetic typology of the $f_{dM}$ analysis; (((P1, P2), P3), O), where P1, P2 and P3 are populations investigated for introgression and O is the outgroup. The table also denotes which processes the genes are involved in (Process), number of annotated genes assigned to the process (Annotated), number of observed significant genes within the process (Significant), the expected number of significant genes (Expected), and a raw p-value. Only processes with a raw p-value of less than 1 are included in the table.

| **(((North American B TA, North American A TA), Canadian TF), *T. biforme*)** | | | | |
|---|---|---|---|---|
| **Process** | **Annotated** | **Significant** | **Expected** | **Raw p-value** |
| Metabolic process | 2388 | 7 | 6.12 | 0.0012 |
| Pantothenate biosynthetic process | 2 | 1 | 0.01 | 0.0016 |
| Copper ion transport | 6 | 1 | 0.02 | 0.0049 |
| Organic substance metabolic process | 1597 | 4 | 4.09 | 0.0912 |
| **(((North American B TA, North American A TA), Italian TF), *T. biforme*)** | | | | |
| **Process** | **Annotated** | **Significant** | **Expected** | **Raw p-value** |
| Metabolic process | 2388 | 11 | 7.83 | 0.00058 |
| Protein phosphorylation | 303 | 3 | 0.99 | 0.00672 |
| Copper ion transport | 6 | 1 | 0.02 | 0.00797 |
| Pseudouridine synthesis | 7 | 1 | 0.02 | 0.00929 |
| Carbohydrate metabolic process | 213 | 2 | 0.7 | 0.03156 |
| Biological_process | 3020 | 13 | 9.9 | 0.08499 |
| Primary metabolic process | 1512 | 7 | 4.96 | 0.09572 |

TA = *Trichaptum abietinum* and TF = *T. fuscoviolaceum*

**Table S5. Estimated model parameters.** The table shows estimated parameters for the different models tested with *fastsimcoal2* (Excoffier et al. 2021). See Figure S7 for illustrations of the models. The headers (except for the model names) are parameters estimated in the analysis. Parameters for the best model is highlighted in light grey. See the GitHub page for example of parameter estimation and template files (*GitHub address will be added*).

| Model name | Npop0 | Npop1 | Npop2 | Npop3 | Npop4 | Npop5 | Tbw1 | Tbw2 | Tbw3 | Tbw4 |
|---|---|---|---|---|---|---|---|---|---|---|
| nomig | 68673 | 166871 | 68367 | 106745 | NA | NA | 205058 | 91189 | NA | NA |
| mig_ancient | 153841 | 130988 | 60260 | 83895 | NA | NA | 365698 | 322921 | NA | NA |
| mig_2ancient | 175064 | 184860 | 69203 | 109285 | NA | NA | 241285 | 81014 | NA | NA |
| mig_3ancient | 243968 | 321735 | 202473 | 161661 | NA | NA | 2076408 | 2149569 | NA | NA |
| mig_recent | 151618 | 185172 | 57426 | 103008 | NA | NA | 233029 | 49995 | NA | NA |
| mig_2recent | 249507 | 94388 | 68676 | 75408 | NA | NA | 212338 | 1765 | 2453 | NA |
| mig_3recent | 151271 | 176916 | 69615 | 101860 | NA | NA | 191882 | 68860 | NA | NA |
| mig_4recent | 137582 | 92899 | 61313 | 116092 | NA | NA | 1335026 | 1242192 | 75564 | NA |
| mig_ghost | 147810 | 117770 | 62165 | 70301 | 66357 | NA | 27347 | 3802 | 3980 | 4082 |
| mig_2ghost | 203693 | 139057 | 85727 | 91339 | 31008 | NA | 192683 | 4625 | 5820 | 12291 |
| mig_3ghost | 99441 | 199627 | 44307 | 95489 | 8697 | NA | 162766 | 4127 | 1416 | 81597 |
| mig_4ghost | 174489 | 253190 | 72062 | 67152 | 48214 | 68623 | 106757 | 2104 | 3613 | 33568 |
| mig_5ghost | 221459 | 152231 | 100595 | 96448 | 60168 | NA | 212413 | 6967 | 6943 | 4007 |
| mig_6ghost | 98582 | 239473 | 36571 | 114641 | 7912 | NA | 128055 | 2285 | 2473 | 2885 |
| mig_7ghost | 193661 | 144624 | 85059 | 87238 | 62942 | 67945 | 20960 | 4950 | 3950 | 4421 |
| mig_8ghost | 162493 | 202415 | 28410 | 10958 | 1809 | 42420 | 1328373 | 22786 | 882 | 25308 |
| mig_9ghost | 112995 | 215613 | 44149 | 112356 | 8660 | NA | 423851 | 1521 | 4565 | 2276 |

| Model name | Tbw5 | Tbw6 | Tbw7 | Resize1 | Resize2 | Resize3 | Resize4 | Resize5 | mig1 | mig2 |
|---|---|---|---|---|---|---|---|---|---|---|
| nomig | NA | NA | NA | 3.7842780 | 1.1040148 | 3.6354008 | NA | NA | NA | NA |
| mig_ancient | NA | NA | NA | 16.4110324 | 0.5678387 | 1.4124021 | NA | NA | 4.99E-06 | NA |
| mig_2ancient | NA | NA | NA | 3.6690522 | 1.1947390 | 7.8242162 | NA | NA | 8.61E-08 | NA |
| mig_3ancient | NA | NA | NA | 0.1919287 | 1.1519895 | 0.9203845 | NA | NA | 6.32E-06 | NA |
| mig_recent | NA | NA | NA | 3.3260382 | 1.3913197 | 20.1069646 | NA | NA | 1.18E-07 | NA |
| mig_2recent | NA | NA | NA | 8.4229324 | 0.4118750 | 51.4254188 | NA | NA | 6.53E-04 | NA |
| mig_3recent | NA | NA | NA | 4.1196068 | 1.1041883 | 2.8093411 | NA | NA | 8.65E-08 | NA |

| | | | | | | | | | | |
|---|---|---|---|---|---|---|---|---|---|---|
| mig_4recent | NA | NA | NA | 1.0985346 | 0.2360949 | 4.3962013 | NA | NA | 0.0078392 | NA |
| mig_ghost | NA | NA | NA | 51.8148929 | 0.1368533 | 32.3027957 | 0.1561456 | NA | 6.52E-04 | NA |
| mig_2ghost | NA | NA | NA | 4.4920900 | 0.8896129 | 7.1859076 | 25.4961475 | NA | 5.25E-08 | NA |
| mig_3ghost | NA | NA | NA | 6.0686313 | 1.0769703 | 0.9589982 | 30.4487050 | NA | 1.63E-04 | NA |
| mig_4ghost | 34844 | NA | NA | 7.7386680 | 0.6000751 | 0.2393338 | 16.1633557 | 0.1060555 | 7.51E-06 | NA |
| mig_5ghost | NA | NA | NA | 4.2221569 | 0.9345088 | 0.8658364 | 45.1669535 | NA | 8.66E-08 | NA |
| mig_6ghost | 71762 | NA | NA | 6.1241827 | 0.8451170 | 39.8374047 | 0.1301317 | NA | 1.77E-05 | 2.12E-04 |
| mig_7ghost | 3388 | NA | NA | 38.3739585 | 0.1179344 | 2.8160925 | 0.1217412 | 6.2096511 | 4.42E-06 | NA |
| mig_8ghost | 1676 | 6450 | 0 | 2.7466486 | 20.3338345 | 0.1026876 | 15.1573729 | 3.7116024 | 2.29E-04 | 6.57E-05 |
| mig_9ghost | 86633 | NA | NA | 15.9185133 | 0.6420154 | 63.8036232 | 0.4883565 | NA | 1.48E-04 | 1.42E-06 |

| **Model name** | **mig3** | **prop1** | **prop2** | **prop3** | **prop4** | **prop5** | **prop6** | **Tdiv1** | **Tdiv2** | **TdivG1** | **Tdiv3** | **TdivG2** |
|---|---|---|---|---|---|---|---|---|---|---|---|---|
| nomig | NA | NA | NA | NA | NA | NA | NA | 354864 | 149806 | NA | 263675 | NA |
| mig_ancient | NA | 1.3078478 | NA | NA | NA | NA | NA | 508998 | 143300 | NA | 186077 | NA |
| mig_2ancient | NA | 0.0292000 | NA | NA | NA | NA | NA | 397096 | 155811 | NA | 316082 | NA |
| mig_3ancient | NA | 1.1130978 | NA | NA | NA | NA | NA | 2430173 | 353765 | NA | 280604 | NA |
| mig_recent | NA | 0.0226856 | NA | NA | NA | NA | NA | 369159 | 136130 | NA | 319164 | NA |
| mig_2recent | NA | 0.0104990 | 0.0147415 | NA | NA | NA | NA | 380536 | 168198 | NA | 166433 | NA |
| mig_3recent | NA | 0.0150532 | NA | NA | NA | NA | NA | 339611 | 147729 | NA | 270751 | NA |
| mig_4recent | NA | 0.3266287 | NA | NA | NA | NA | NA | 1473540 | 138514 | NA | 231348 | NA |
| mig_ghost | NA | 0.0251477 | 0.0270060 | 0.0284662 | NA | NA | NA | 178536 | 151189 | 147387 | 139325 | NA |
| mig_2ghost | NA | 0.0215626 | 0.0277299 | 0.0602280 | NA | NA | NA | 407206 | 214523 | 209898 | 204078 | NA |
| mig_3ghost | NA | 0.0440787 | 0.0158260 | 0.9262883 | NA | NA | NA | 256400 | 93634 | 89507 | 88091 | NA |
| mig_4ghost | NA | 0.0135652 | 0.0236086 | 0.2246313 | 0.3007132 | NA | NA | 261914 | 155157 | 153053 | 149440 | 115872 |
| mig_5ghost | NA | 0.0294961 | 0.0302867 | 0.0180291 | NA | NA | NA | 448629 | 236216 | 229249 | 218299 | NA |
| mig_6ghost | NA | 0.0278678 | 0.0310197 | 0.0373450 | 0.9648624 | NA | NA | 210074 | 82019 | 79734 | 77261 | NA |
| mig_7ghost | NA | 0.0247137 | 0.0202190 | 0.0230985 | 0.0181224 | NA | NA | 221273 | 200313 | 195363 | 183604 | 191413 |
| mig_8ghost | 3.22E-04 | 0.3068627 | 0.0171475 | 0.5003000 | 0.0663087 | 0.2733081 | 0.0175051 | 1402628 | 74255 | 51469 | 23603 | 50587 |
| mig_9ghost | NA | 0.0151238 | 0.0460690 | 0.0240852 | 0.9390135 | NA | NA | 524473 | 99101 | 94536 | 92260 | NA |

| Model name | Tmig1 | Tmig2 | Tmig3 | MaxEstLhood | MaxObsLhoos | AIC |
|---|---|---|---|---|---|---|
| nomig | NA | NA | NA | -7823474.701 | -6809027.340 | 1014447.361 |
| mig_ancient | 187414 | NA | NA | -7805505.703 | -6809510.928 | 995994.775 |
| mig_2ancient | 4549 | NA | NA | -7819058.784 | -6809510.928 | 1009547.856 |
| mig_3ancient | 312339 | NA | NA | -7792802.746 | -6809510.928 | 983291.818 |
| mig_recent | 3088 | NA | NA | -7821830.565 | -6809510.928 | 1012319.637 |
| mig_2recent | 163980 | NA | NA | -7764526.353 | -6809510.928 | 955015.425 |
| mig_3recent | 4075 | NA | NA | -7813832.196 | -6809510.928 | 1004321.268 |
| mig_4recent | 155784 | NA | NA | -7728427.346 | -6809510.928 | 918916.418 |
| mig_ghost | 143407 | NA | NA | -7808874.997 | -6809510.928 | 999364.069 |
| mig_2ghost | 191787 | NA | NA | -7845453.172 | -6809510.928 | 1035942.244 |
| mig_3ghost | 6494 | NA | NA | -7664400.865 | -6809510.928 | 854889.937 |
| mig_4ghost | 81028 | NA | NA | -7791242.284 | -6809510.928 | 981731.356 |
| mig_5ghost | 222306 | NA | NA | -7845664.969 | -6809510.928 | 1036154.041 |
| mig_6ghost | 74376 | 2614 | NA | -7650743.782 | -6809510.928 | 841232.854 |
| mig_7ghost | 186992 | NA | NA | -7821700.972 | -6809510.928 | 1012190.044 |
| mig_8ghost | 25279 | 17153 | 17153 | -7754011.360 | -6809510.928 | 944500.432 |
| mig_9ghost | 100622 | 5627 | NA | -7644407.431 | -6809510.928 | 834896.503 |
